# Discovering tumor-reactive T-cell receptors through single-cell sequencing of tumor-infiltrating lymphocytes

**DOI:** 10.1101/2021.11.30.470597

**Authors:** Elinor Gottschalk, Bulent Arman Aksoy, Pinar Aksoy, Marzena Swiderska-syn, Caroline Mart, Linda Macpherson, Martin Romeo, Aubrey Smith, Hannah Knochelmann, Chrystal Paulos, Cynthia Timmers, John Wrangle, Mark P. Rubinstein, Jeff Hammerbacher

## Abstract

We evaluated the utility of single-cell sequencing of tumor-infiltrating lymphocytes (TIL) for tumor-reactive T-cell receptor (TCR) discovery. Using the MC38 cell line as our tumor model in mice, we show that expression of exogenous TCRs via mRNA electroporation in human T cells provides an easy and quick path to validating tumor-specific candidate TCRs. We detail the identification and validation of four novel MC38-reactive mouse TCRs with varying levels of reactivity to the target cells. Validating our process, one of the MC38 TCRs is specific against a previously reported neoantigen (ASMTNMELM in the Adpgk gene). Consideration of these methodologies may aid in the development of rapid TCR-based therapies for the treatment of cancer and human disease.

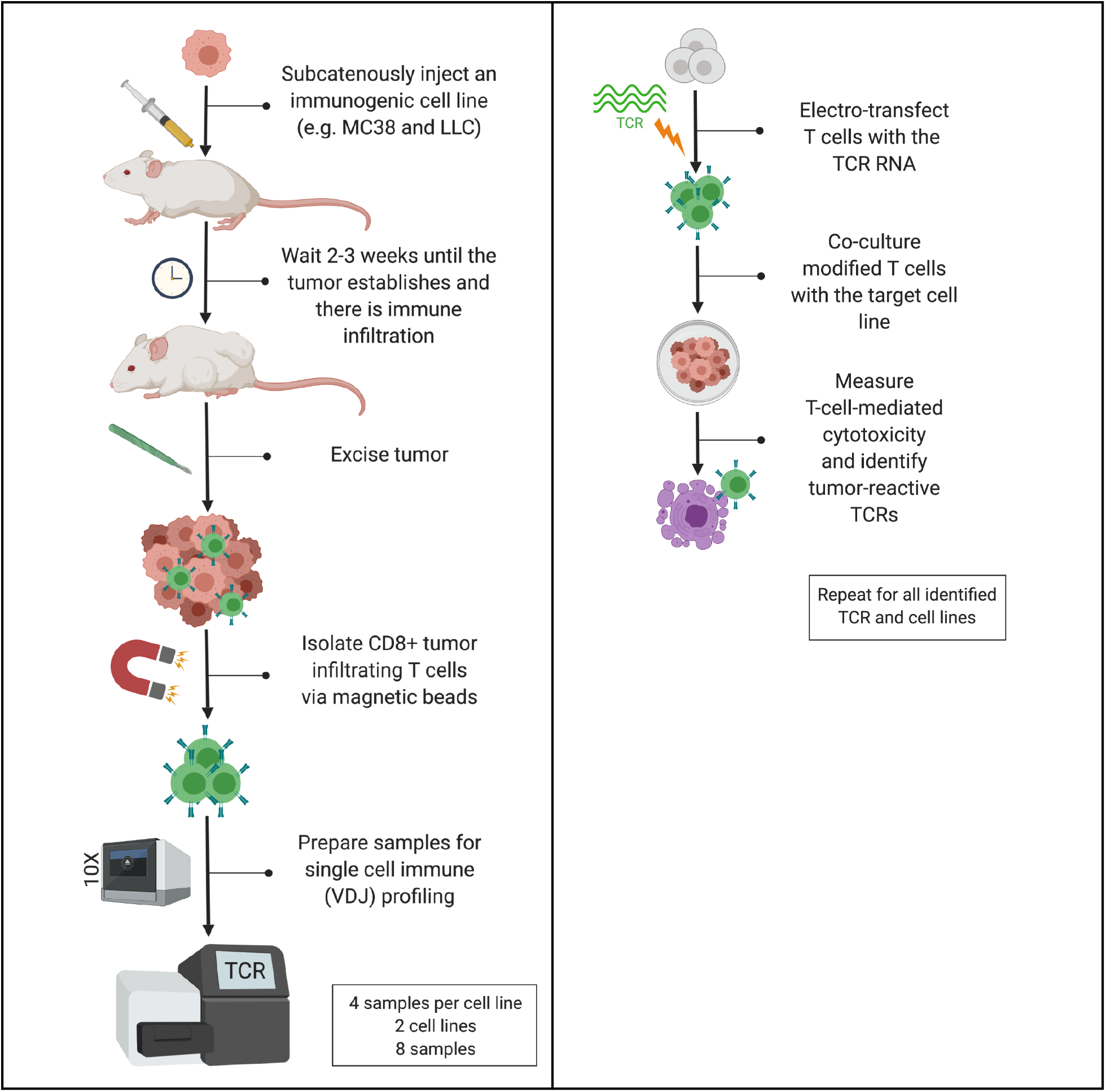

## Introduction

Tumor-infiltrating lymphocytes (TILs) are a valuable resource for adoptive cell therapy (ACT) as one immunotherapeutic approach to treat cancer. TILs have a high chance of being tumor-reactive, meaning they can recognize tumor cells from their parent tumor and eliminate them. The method of ACT pioneered by Rosenberg takes advantage of a patient’s TILs for personalized cell therapy by expanding the cells from tumor samples *ex vivo* and, when the cell numbers are sufficient, transferring the expanded cell product to patients via infusion (Rosenberg and Restifo 2015).

The alpha and beta T cell-receptor (TCR) genes provide a T cell with the ability to recognize a specific antigen (Cole et al. 1995). TILs, which are often T cells, recognize tumor antigens on a tumor cell and are considered tumor-reactive. The expansion of TILs is essentially cloning T cells with tumor-reactive TCRs. Identification of TCRs from TILs often requires that TILs be expanded to adequate numbers. TILs profiled directly after isolation from the tumor are often found in an exhausted state, which limits their proliferative capacity and can negatively affect their effector function (Miller et al. 2019; Baitsch et al. 2011; Ahmadzadeh et al. 2009; Wherry 2011; Blank et al. 2019). For any TILs that are not already exhausted from residing in the tumor, the long *ex vivo* expansion in high IL-2 conditions to obtain sufficient cell numbers for ACT can push the cells into an exhausted state. Furthermore, it is unclear how the *ex vivo* expansion of TILs affects the initial diversity of TCRs in a TIL population.

While TILs are one source for cells to use in ACT, the other major approach is to isolate T cells from a patient’s peripheral blood and engineer the cells to express TCRs or chimeric antigen receptors (CARs) to target a known antigen on tumor cells (Met et al. 2019). The main limitation of this approach is the requirement of known TCR or CAR sequences and their targets.

Here we extracted sequences of naturally-arising TCRs in TILs, which were then used to engineer non-exhausted T cells. Our goal was to test the approach in a mouse tumor model and determine the feasibility of quickly obtaining tumor-reactive TIL-derived TCRs for subsequent expression. The technologies to sequence TCRs (also known as, immune or V(D)J profiling) on a single-cell level and the ability to genetically manipulate human T cells have been challenging and infeasible processes so far due to technological limitations. However, we now have affordable access to reliable single-cell sequencing (to obtain paired TCR sequences on a per cell basis), we know how to efficiently manipulate human T cells with arbitrary genetic material, and we have adapted a reliable method for measuring TCR reactivity. Here, we show that it was possible to quickly obtain a pool of tumor-reactive TCR sequences from TILs directly after isolation from an MC38 tumor, that there was an enrichment of a different TCR in each tumor, and that the most prominent TCR from each MC38 tumor was tumor-reactive. Additionally, we successfully de-orphanized one of the four MC38 TCRs that we investigated, identifying the Adpgk-derived peptide ASMTNMELM as the cognate antigen. Our investigation with an alternate cell line, LLC-A9F1, similarly resulted in enriched TCR sequences from each tumor’s TIL population, however, we did not complete the validation of these TCR’s tumor-reactivity for this preprint.

## Results

### V(D)J profiling of freshly isolated TILs showed an enrichment of one TCR clonotype in each tumor

As human tumor samples are heterogeneous, scarce, and a finite resource, we started by testing our idea in an animal model. Mouse models have the benefits of consistency (little variation from animal to animal), on-demand access to tumor samples, and the capability to induce tumors from a cell line (which is an infinite resource). Our immediate goal was to subcutaneously inject a cell line into a mouse; let it establish as a tumor and attract immune cells; excise the tumor and profile TCRs from TILs; express the new TCR pairs in T cells (one TCR at a time); and assess T-cell mediated cytotoxicity of these transformed T cells on the cancer line. We used two cell lines for this study: MC38, a mouse colon adenocarcinoma, and LLC-A9F1, a Lewis lung mouse carcinoma. Both MC38 and LLC-A9F1 are immunogenic tumors, thus increasing the chance of tumor-reactive TILs being present in the tumor for us to isolate and sequence (Eisenbach, Segal, and Feldman 1983; Sultan et al. 2020; Hos et al. 2019; Yadav et al. 2014).

Our first goal was to test the feasibility of obtaining paired tumor-reactive TCRs from TILs directly after isolation from tumors. A total of 20 female C57BL/6 (H-2K^b^/H-2D^b^) mice were subcutaneously injected with either MC38 (10 mice) or LLC-A9F1 (10 mice) tumor cells. The mice were housed 5 mice per cage and 2 cages per tumor cell line. The tumors were allowed to grow until the largest was about 100 mm^3^ before being excised (tumor growth data is linked in the Materials and Methods). The largest 8 tumors from each cell line were subsequently processed. TILs were immediately isolated after excision, after which CD8+ T cells were positively isolated with Dynabeads. Directly after isolation, the CD8+ T cells were assessed for viability and the highest quality samples were prepared for scTCR sequencing. From the 8 tumors for each cell line, 4 from each were at a high enough quality (above 60% viability) to proceed with single cell sequencing. The CD8+ T cells from each sample were submitted to the MUSC Translational Core, where the samples were prepared with the 10x V(D)J profiling kit (further details on the sample preparation is provided in materials and methods) and submitted to Vanderbilt University for sequencing. 3000-4000 cells from each tumor sample were sequenced. The cell counts for each sample are shown in Table 1, along with the total barcodes and clonotypes found per tumor sample. The total number of barcodes is lower than the cell count because only barcodes with productive CDR3s can be included in a clonotype. A clonotype is defined as the unique set of complementarity-determining region 3s (CDR3s) in a cell. CDR3s can be shared between cells to make different clonotypes (i.e. two cells can express the same α TCR subunit but different β subunits). Approximately 700 - 1500 TCR clonotypes were identified in each tumor sample. However, not all of the clonotypes identified consist only of a single α and single β chain (some clonotypes contain only one chain total and some contain more than one α or β chain), which is consistent with the literature on allelic inclusion in T cells (Malissen et al. 1992; Brady, Steinel, and Bassing 2010).

**Table 1.** Cell counts, total number of barcodes, and clonotypes for the TILs isolated from each tumor sample. The TILs from each tumor were prepared and sent for single cell immune profiling with the 10x V(D)J immune profiling kit. The cell line and tumor number for each sample is listed and the TCR that was subsequently investigated from each tumor sample is stated in parentheses. Data were analyzed in the 10x Genomics Loupe V(D)J Browser.

| Sample | Cell Count | Total barcodes | Clonotypes |
| --- | --- | --- | --- |
| MC38 tumor 1 (TCR30) | 3985 | 3950 | 1333 |
| MC38 tumor 2 (TCR31) | 3382 | 3363 | 1144 |
| MC38 tumor 3 (TCR32) | 4347 | 4303 | 1098 |
| MC38 tumor 4 (TCR33) | 3887 | 3834 | 808 |
| LLC-A9F1 tumor 1 (TCR34) | 3304 | 3258 | 1574 |
| LLC-A9F1 tumor 2 (TCR35-1 and 35-2) | 3047 | 3013 | 1464 |
| LLC-A9F1 tumor 3 (TCR36-1 and 36-2) | 2684 | 2672 | 728 |
| LLC-A9F1 tumor 4 (TCR37) | 3070 | 3047 | 816 |

The top 10 most frequent clonotypes that were identified in each sample are shown in Figure 1 as percent of total barcodes (the barcode frequency for each clonotype is listed on the x-axis). There was an enrichment of one clonotype in the majority of the samples. Each MC38 sample had one TCR pair that was the most abundant (being present in 20 - 40% of the sequenced TILs). The LLC-A9F1 results showed more variation, with the main clonotypes in two tumors being present in approximately 10% of the TILs and in 40-50% of the TILs in the other two tumors. The top clonotype in the LLC-A9F1 tumor 2 had two α TCR chains, which we split into two TCR combinations for testing (35-1 and 35-2). Likewise, the top clonotype in LLC-A9F1 tumor 3 had two β chains, which we separated for testing with the α chain (36-1 and 36-2). The top clonotype in LLC-A9F1 tumor 4 was identical to the top clonotype in tumor 3 (it contained the same α chain and same two β chains).

**Figure 1.**
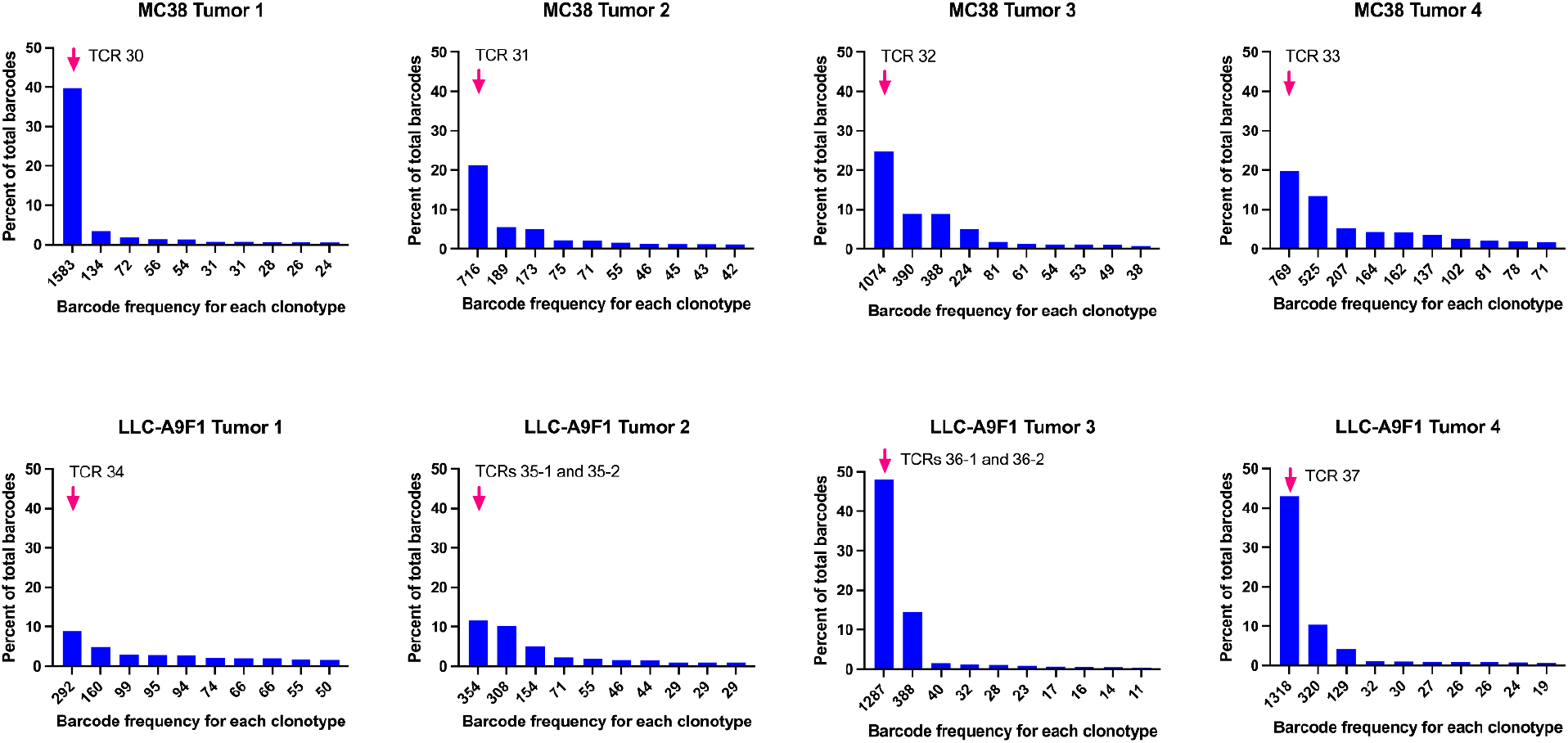
Direct single cell sequencing of TILs reveals one prominent clonotype per tumor. The top 10 most frequent clonotypes per tumor sample are shown. The percent of each clonotype per total barcodes is shown on the y-axis. The number of cells with each clonotype is listed on the x-axis. The clonotype from each tumor chosen for functional analysis is indicated by a pink arrow and the initial name of the TCR pair/s is listed. Data were analyzed in the 10x Genomics Loupe V(D)J Browser and graphs were created with GraphPad Prism 9.

The number of shared clonotypes in the TILs between the four MC38 tumor samples was limited (Figure 2, left panel). Tumor 1 (sample 30) shared 1 clonotype with tumors 2 (31) and 3 (32), and 2 clonotypes with tumor 4 (33). Tumor 2 (31) did not share any other clonotypes with TILs from the other two tumors (only the one shared clonotype with 30). Tumor 3 (32) shared 11 clonotypes with 33, in addition to the one with tumor 1 (30). There was a higher number of shared clonotypes between the LLC-A9F1 samples (Figure 2, right panel). The lowest being 2 shared clonotypes between two samples, and the highest being 38 between tumor 3 (36) and 4 (37).

**Figure 2.**
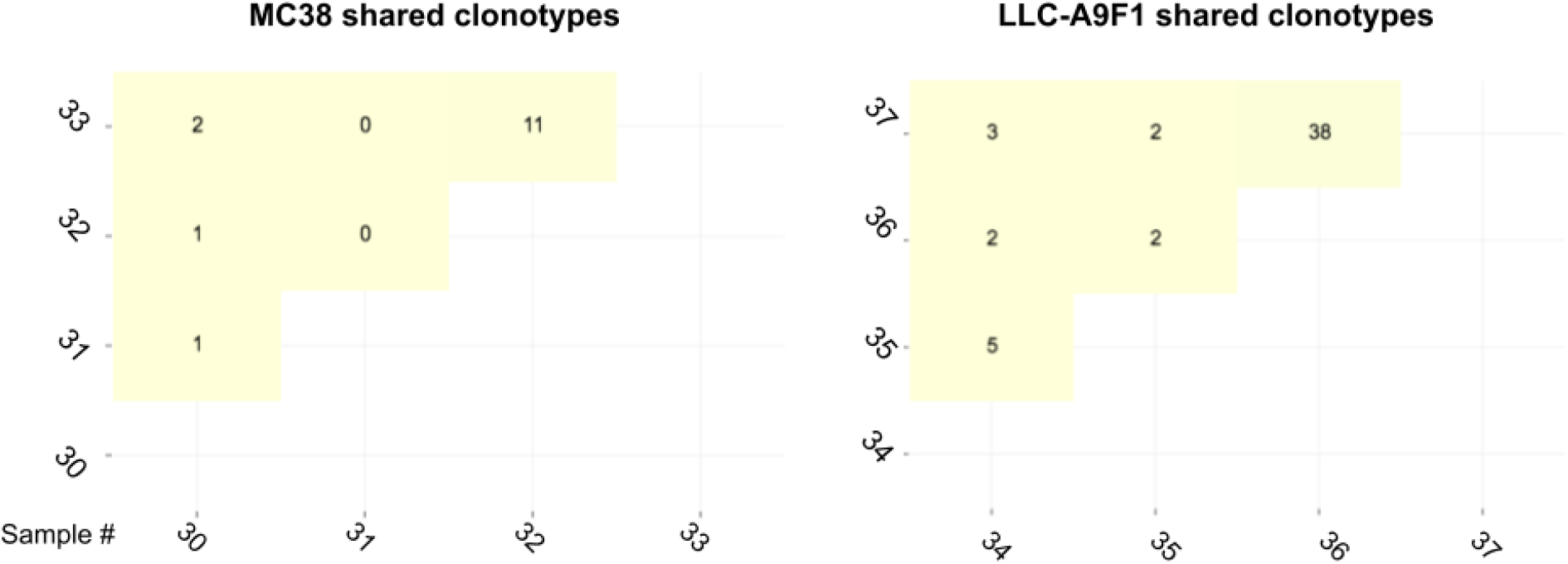
Shared clonotypes between the four MC38 tumors and the four LLC-A9F1 tumors. Data were analyzed in the 10x Genomics Loupe V(D)J Browser.

We were curious whether the most frequent clonotypes are the ones that are shared between samples. Interestingly, the one clonotype that tumor sample 32 shared with 30 is the most prominent clonotype in sample 30 (TCR30, which is the TCR clonotype we chose to investigate from this tumor), but was at a very low frequency in sample 32. This clonotype was in approximately 40% of the TILs in sample 30 and only 0.02% in sample 32. This clonotype was not detected in tumor samples 31 and 33. TCR 30 is the only shared clonotype from sample 30 that has both an α and β chain. The other shared clonotypes only had one β chain and were at a prevalence of less than 1%. The one clonotype that sample 31 shares with 30 was at low frequency in both samples (0.03% and 0.06%, respectively). The two clonotypes shared between sample 30 and 33 were also at low frequencies for both samples (between 0.03% and 0.4%). Samples 32 and 33 shared 11 clonotypes, the most out of the MC38 samples. The most frequent clonotype in sample 32 (at 24%) was present at 0.2% in sample 33. MC38 shared clonotypes that were present in at least 1% of the TILs from each tumor are summarized below in Table 2, including the CDR3 amino acid sequences for the α and β chains.

**Table 2.** Clonotypes shared between the four MC38 tumor samples (30-33). The TCR chains making up the clonotypes and their CDR3 amino acid sequences are listed. The percentage that each clonotype occupies of the total barcodes in each sample is shown. Only the clones with at least 1% prevalence in any of the tumor samples were listed.

| TCR chain | CDR3 amino acid sequences | 30 | 31 | 32 | 33 |
| --- | --- | --- | --- | --- | --- |
| TRA;TRB | CAIDPYNQGKLIF;CASSPGQGYAEQFF | 39.71% | 0.00% | 0.02% | 0.00% |
| TRA;TRB | CAVSDDSNYQLIW;CASSRDNSYNSPLYF | 0.00% | 0.00% | 24.71% | 0.26% |
| TRB | CASSRDNSYNSPLYF | 0.00% | 0.00% | 8.93% | 0.05% |
| TRB | CGAWTVSNERLFF | 0.00% | 0.00% | 1.86% | 0.03% |

The most frequent TCR clonotype found in each tumor contained one at least one of each an α and β TCR chain, which allowed us to test αβ pairs *in vitro*. The full TCR sequences were constructed for cloning and expression as described in the Materials and Methods.

### MC38 tumor TIL-derived TCRs can kill MC38 cells in co-culture

We began our investigation of the newly identified TCRs by focusing on the MC38 samples. We wanted to test whether the MC38 TCRs had the ability to recognize and kill MC38 cells (their parental tumor) in co-culture. We chose the most frequent TCR pair from each sample, with the assumption that the top clonotype would be the most likely to be tumor-reactive. We conducted cytotoxicity assays by expressing the MC38 TCRs in T cells, via mRNA electroporation, and co-culturing the modified T cells with MC38 cells. In our lab’s previous work we optimized the expression of OT-I TCRs in primary mouse or human CD8+ cells via mRNA electroporation and showed that the modified cells had the ability to recognize and kill SIINFEKL-presenting cells using a flow cytometry-based assay (Aksoy et al. 2019). Here, we started with the same flow cytometry-based assay to test the MC38 TCRs.

We first attempted to express the MC38 TCRs in activated mouse CD8+ T cells isolated from splenocytes. However, we experienced a very low efficiency with the OT-I TCR when we stained the samples with an OT-I dextramer (data not shown). Therefore, we moved to using CD8+ T cells isolated from healthy human donors. Briefly, we expressed either the OT-I TCR (as a control) or the MC38 TCRs in activated CD8+ T cells isolated from 3 healthy human donors (via mRNA electroporation) and co-cultured them with MC38 cells for 24 hours. All of the cells were harvested (T cells and cancer cells) and prepared for flow cytometry. The samples were stained with CD3 and CD8 antibodies in order to gate out the T cell population during analysis, which allowed the remaining live MC38 cells to be counted in each sample. The full protocol for this assay is linked in materials and methods. In our assay we used OT-I TCR and its cognate antigen (SIINFEKL) as controls. Conditions without the SIINFEKL peptide are the negative control (no MC38 killing) and the addition of the peptide is the positive control. Figure 3, panel A shows the MC38 cell counts for each control condition with T cells from the three human donors (black bars show the samples that had SIINFEKL peptide; pink bars did not have the peptide added). There was a marked decrease in MC38 cell number when SIINFEKL was included in the cultures with the OT-I TCR-expressing T cells. Panel B in Figure 3 shows what the percent cytotoxicity was with the 3 donors, using the cell counts shown in panel A. In Panel C, our results showed that, when simply co-cultured with MC38 cells, T cells expressing each of the four novel MC38 TCRs were successful in killing between 50 - 95% of the cancer cells after normalization to mock-electroporated donor T cells (which were not engineered to express an exogenous MC38 TCR).

**Figure 3.**
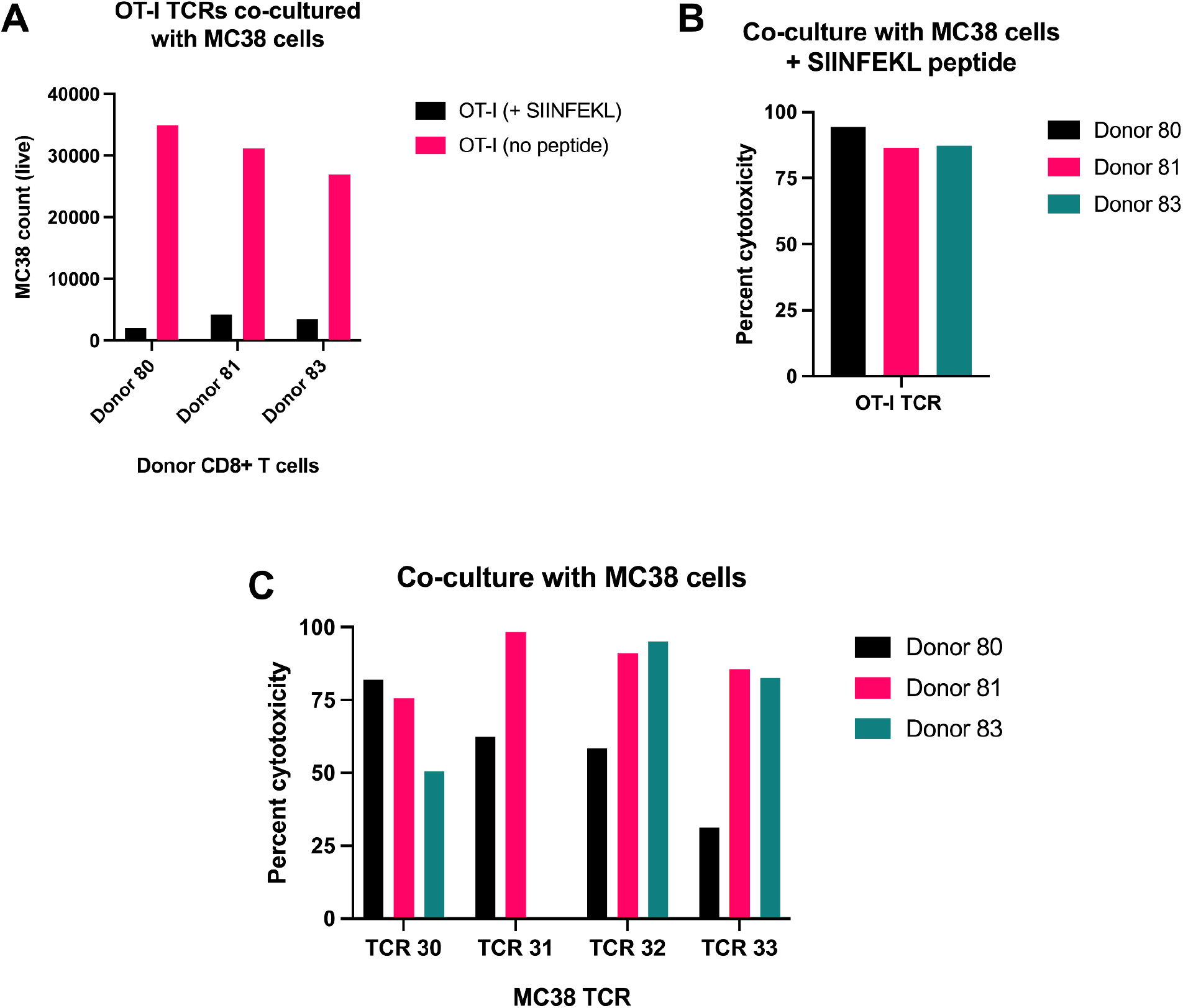
Human T cells expressing murine TCRs kill MC38 cancer cells in co-culture. Results from a flow cytometry-based cytotoxicity assay with OT-I TCR (control) or MC38 TCRs are shown. CD8+ T cells were isolated from three healthy human donors (Human donors 80, 81, and 83). TCR mRNA was electroporated into human CD8+ T cells and co-cultured with MC38 cancer cells overnight at an 8:1 T cell:cancer cell ratio. A) The OT-I TCR was used as a control. Co-cultures with the addition of the cognate peptide (SIINFEKL) as a positive control (black bars) and co-cultures without the peptide as a negative control (pink bars). Live MC38 cell counts are shown. B) Percent cytotoxicity was calculated for each donor expressing the OT-I TCR with SIINFEKL peptide added to the co-culture. C) The percent cytotoxicity was calculated for MC38 cells co-cultured with T cells from each donor expressing the four MC38 TCRs after normalizing to a mock-electroporated control for each donor. The sample from donor 83 for TCR 31 was lost during flow cytometry analysis.

### MC38 tumor TIL-derived TCRs can induce T cell activation in co-culture with MC38 cells

We found limitations to the flow cytometry-based cytotoxicity assay. The number of electroporated T cells we could obtain at one time limited the number of conditions for co-culture, preventing us from varying the ratio of T cells to cancer cells. Additionally, the time required to run each sample to completion on the flow cytometer was significant, which further made it infeasible to expand the experiments to include multiple conditions or subsequent screening of the TCRs for their cognate antigens. Therefore, we explored other options for screening the new TCRs and eventually adapted the Promega T Cell Activation Bioassay Kit for our purposes. The kit contains Jurkat cells that have been modified to express a luciferase reporter upon the activation of Nuclear Factor of Activated T cells (NFAT), which occurs during TCR-mediated activation of T cells. To modify the cells supplied with the kit for our purpose, we needed to be able to co-express the TCR subunits of interest along with the two mouse CD8 subunits in the Jurkat-NFAT cells because Jurkat cells are CD8 negative. To test the co-expression of mCD8 with TCRs, we electroporated the mCD8 subunits along with the OT-I TCR subunits and prepared the samples for flow cytometry 24 hours post-electroporation. We used an OT-I dextramer to check for functional surface expression of the OT-I TCRs (Figure 4A, mock electroporation in grey, mRNA-electroporated in pink). Antibody staining of the alpha and beta subunits of mouse CD8 are shown in Figure 4B (Mock electroporation on the left and mRNA electroporated on the right).

**Figure 4.**
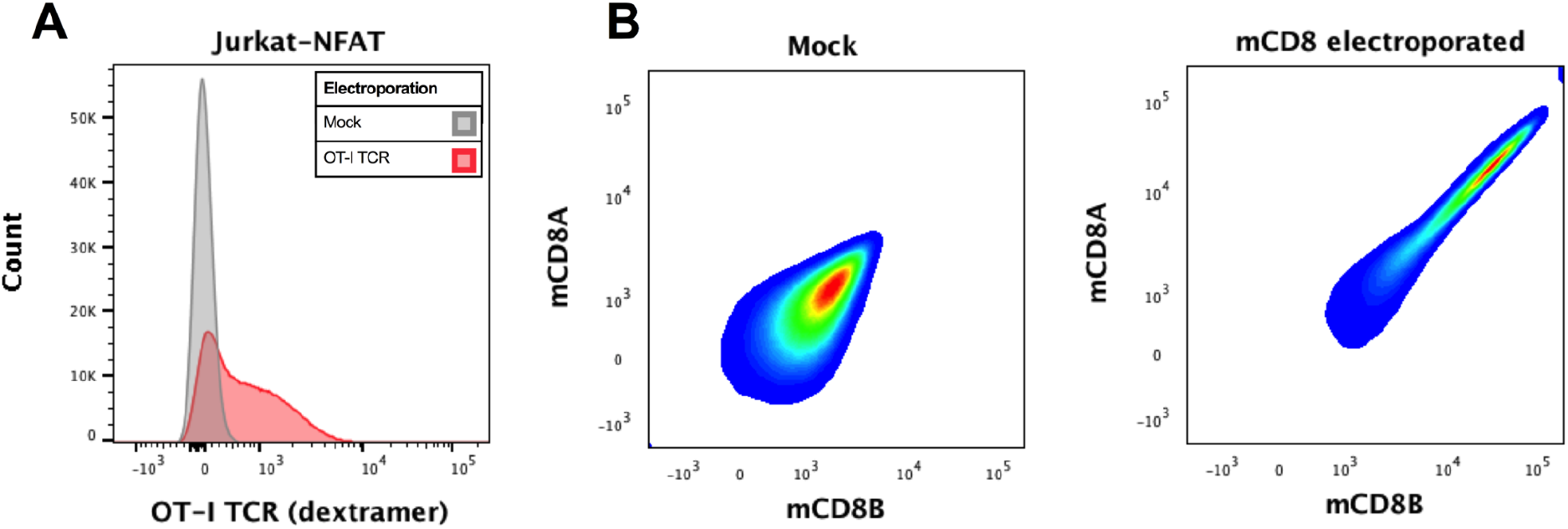
Expression of the OT-I TCR and mCD8 in Jurkat-NFAT cells via mRNA electroporation. Jurkat-NFAT cells were either mock electroporated (no mRNA) or co-electroporated with mRNA encoding the alpha and beta subunits for the OT-I TCR and mouse CD8. The next day the cells were stained with an OT-I dextramer (A) and antibodies against mCDA and mCD8B (B) and analyzed by flow cytometry. n=4.

Our results suggested that the OT-I TCRs were being expressed and functional when electroporated into the Jurkat-NFAT cells along with the mouse CD8 subunits. We moved on to using the Jurkat-NFAT system with the MC38 TCRs, including the OT-I TCR as a control (with/without SIINFEKL peptide). With this assay we were able to expand our range of MC38 cells to T cell ratios from 2:1 to 1:64. We consistently observed high luminescence values with our positive control, OT-I TCR with SIINFEKL (Figure 5, black line) and low levels with the negative control, OT-I TCR and no peptide (Figure 5, pink line). We considered the signal from the negative control condition to be baseline luminescence for comparing any signal from the new TCRs. All four of the MC38 TCRs were capable of activating the Jurkat-NFAT cells when co-cultured with MC38 cells (Figure 5, teal, purple, lavender, and blue lines) and the activation level was dependent on the ratio of T cells to cancer cells; the higher number of cancer cells to T cells, the more T cell activation. Interestingly, the four different MC38 TCRs resulted in varying levels of activation when co-cultured with MC38 cells. We consistently observed that TCR 30 resulted in the lowest activation signal and TCR 32 the highest.

**Figure 5.**
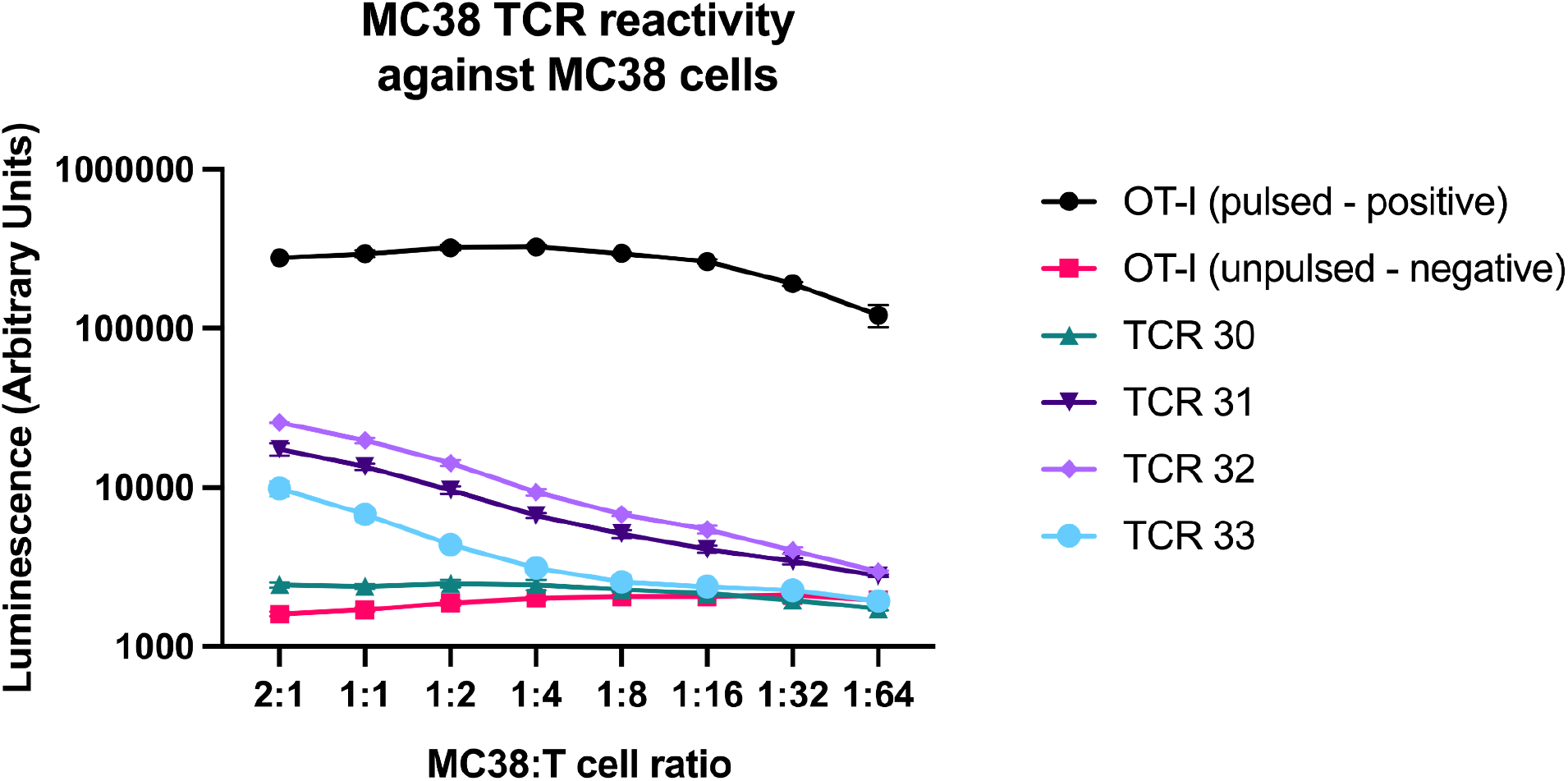
MC38 TCRs result in T cell activation when co-cultured with MC38 cells. Jurkat-NFAT cells expressing either the OT-I TCR or one of the MC38 TCRs were co-cultured with MC38 cells for 12 hours before luciferase expression was measured on a luminescence-based plate reader. The positive control is OT-I pulsed (indicating the addition of the SIINFEKL peptide to the cultures with the Jurkat-NFAT cells expressing the OT-I TCR) and the negative control is OT-I unpulsed, indicating no addition of SIINFEKL peptide (black and pink lines, respectively). N=6 for each condition and the mean and SD from the mean are shown. Representative results from 1 of 5 experiments are shown.

### One MC38 TCR, TCR 30 (Picky sticky), is reactive against previously described neoepitope peptides

Once our data suggested that the four MC38 TCRs were tumor-reactive, we were interested in whether we could find any of their cognate antigens. In our first effort to de-orphanize the MC38 TCRs we took advantage of MC38 neoepitopes that have been described by other investigators. We tested two sets of MC38 neoantigens that have been published, listed in Table 3 (Yadav et al. 2014; Hos et al. 2019). In these experiments we used the modified T Cell Activation Bioassay as described above, with the addition of each neoantigen peptide to the co-cultures and co-cultured for 6 hours before measuring luciferase expression. Our hypothesis was that if a particular MC38 TCR was reactive to a peptide, that there would be an increase in T cell activation (indicated by an increase in luciferase luminescence) beyond the activation we observed when co-culturing the TCRs with MC38 cells alone.

**Table 3.**
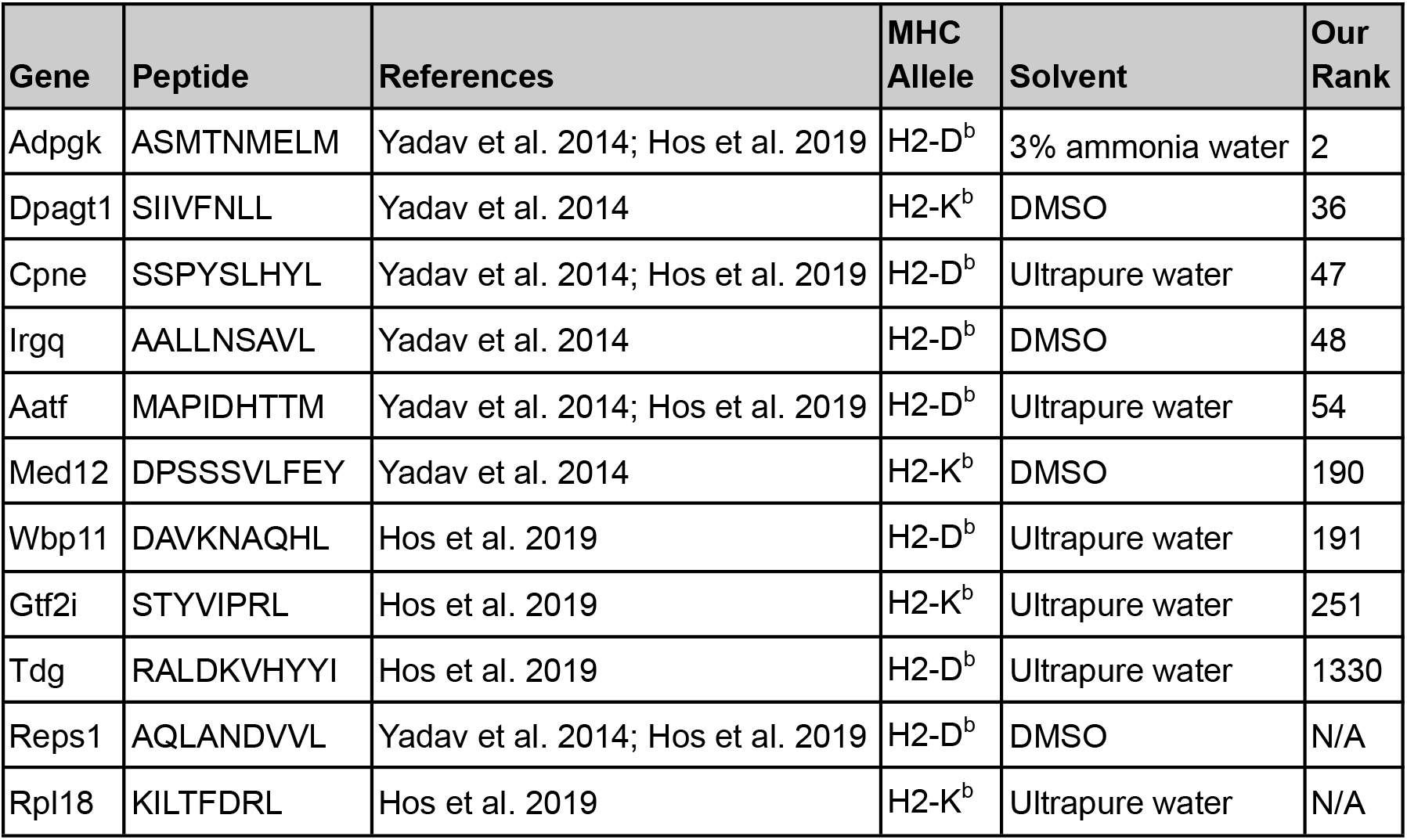
The previously identified MC38 neoepitope peptides first tested with the MC38 TCRs. The name of the gene from which each peptide is derived, the amino acid sequence used in our experiments, references for the neoepitopes, and the predicted MHC allele dependency are listed. The solvent we used to dissolve each peptide is listed. The solvents were recommended by Genscript with their Peptide Solubility Test service. The rank of each peptide in our neoantigen prediction pipeline (described in Methods) is listed in the final column. We ranked peptides with a TPM greater than one based on their predicted affinity. Two peptides were not present in our predictions (Reps1 and Rpl18) because these mutations are not present in our sequencing of the MC38 cell line.

Results from the T cell activation bioassay with the MC38 peptides described above are shown in Figure 6. Again, we expressed the OT-I TCR in the Jurkat-NFAT cells to use as positive and negative controls, with or without the addition of SIINFEKL peptide, respectively (black bars A-D). Each graph shows the T cell activation for one of the MC38 TCRs co-cultured with the peptides listed in Table 3. The horizontal dotted line on each graph indicates the level of activation observed when the TCR is co-cultured with MC38 cells alone. We were looking for an increase from the activation we had seen when the TCRs were co-cultured with MC38 cells alone, as shown in Figure 5. Jurkat-NFAT cells expressing TCR 30 (Picky Sticky) co-cultured with two different peptides (one at a time) derived from the genes Med12 and Adpgk, consistently resulted in increased T cell activation (Figure 6A). A second peptide, derived from the Med12 gene, also consistently increased reactivity of TCR 30 (Figure 6A, last bar). However the increase in activation appeared to be much less than with the Adpgk-derived peptide.

**Figure 6.**
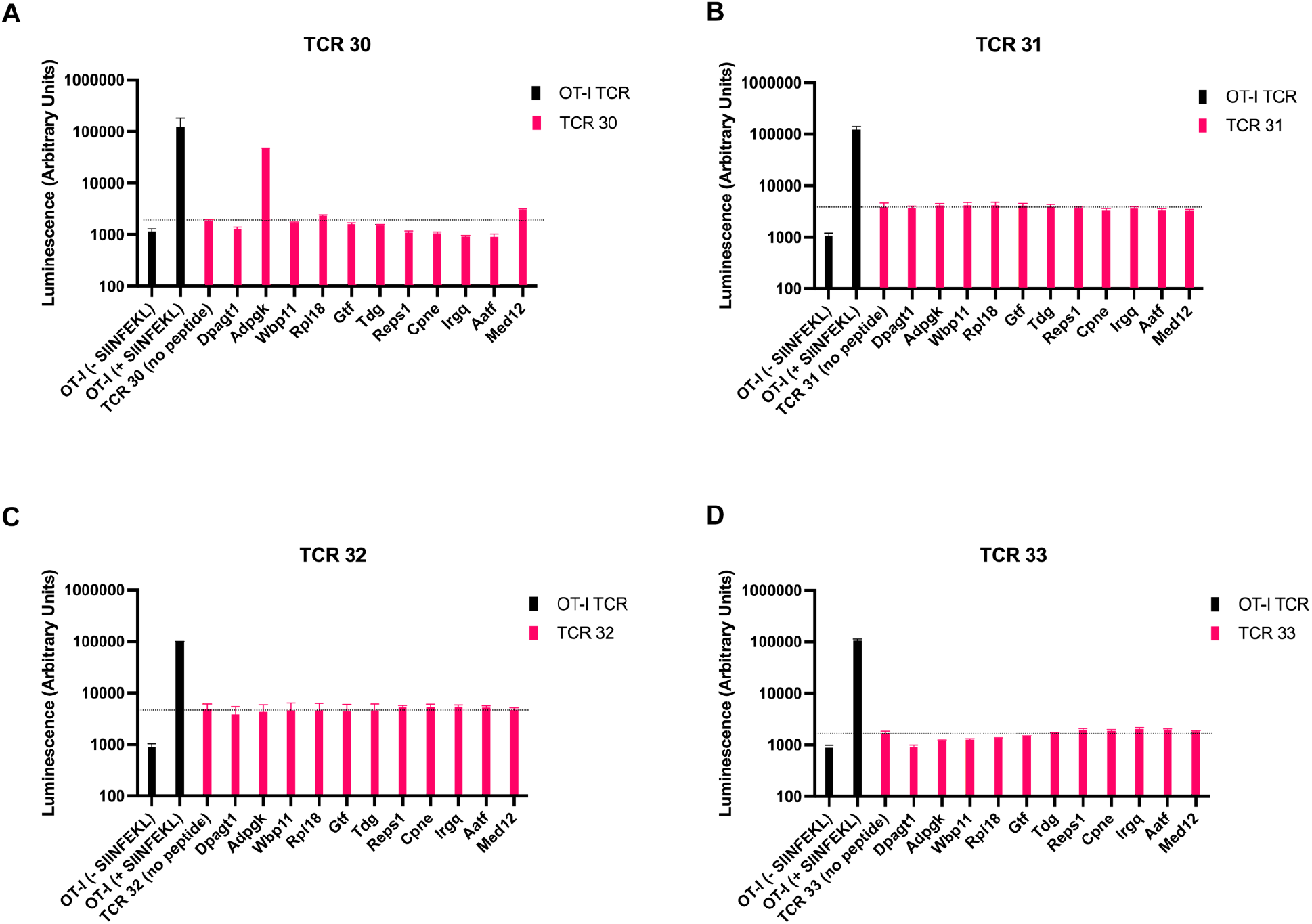
TCR-mediated T cell activation levels of MC38 TCR-expressing cells when co-cultured with predicted neoantigen peptides and MC38 cells. Jurkat-NFAT cells expressing either the OT-I TCR or one of the MC38 TCRs were co-cultured with MC38 cells for 6 hours before being assayed for luciferase expression on a luminescence-capable plate reader. The positive control is OT I pulsed (indicating the addition of the SIINFEKL peptide to the cultures with the Jurkat-NFAT cells expressing the OT-I TCR) and the negative control is OT-I unpulsed, indicating no addition of SIINFEKL peptide (black bars on A-D). Pink bars indicate conditions with one of the MC38 TCRs and an addition of peptide. The dotted black line is drawn at the average activation for each TCR co-cultured with the MC38 cells alone (i.e. no peptide was added). N=6 for each condition and the mean and SD from the mean are shown.

Because TCR 30 was reactive against two of the predicted neoantigen peptides we titrated the amount of each peptide and co-cultured with TCR 30-expressing Jurkat-NFAT cells at a fixed ratio of 2:1 of MC38 cells to T cells (Figure 7). As a positive control we co-cultured OT-I-expressing Jurkat-NFAT cells with MC38 cells and titrated the SIINFEKL peptide added to the cultures (Figure 7, black line). Even at very low concentrations (.01 nM) of SIINFEKL peptide the activation was much higher than the condition with no peptide. As a negative control we co-cultured TCR 30-expressing cells with a titration of SIINFEKL peptide and observed no T cell activation (pink line). The activation levels of TCR 30-expressing T cells with the titration of the Adpgk-derived peptide (purple line) showed a similar curve as the SIINFEKL + OT-I TCR condition. Both SIINFEKL and Adpgk-derived peptides appear to saturate the system at 1 - 0.1 uM. The Med12-derived peptide only resulted in increased levels of T cell activation at the higher concentrations of the peptide (10 and 1 uM).

**Figure 7.**
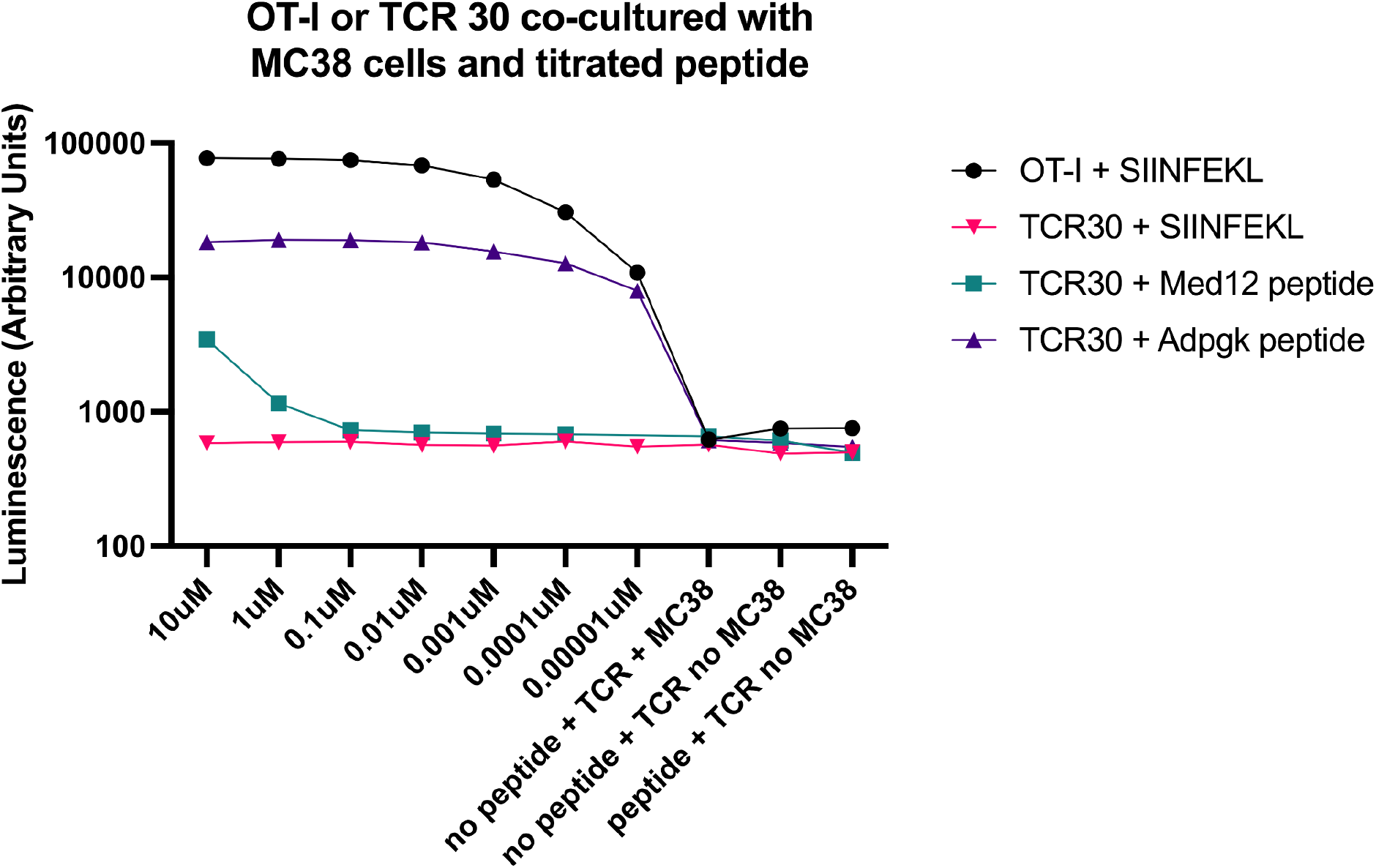
Jurkat-NFAT cells expressing either the OT-I TCR or the MC38 TCR 30 were co-cultured with MC38 cells and a titration of peptide (SIINFEKL, Med12-, or Adpgk-derived peptide). After 6 hours the cultures were assayed for luciferase expression on a luminescence-capable plate reader. The positive control is cells expressing the OT-I TCR with the addition of the SIINFEKL peptide (black line). The pink, teal, and purple lines are cells expressing the MC38 TCR30 with the addition of the SIINFEKL, Med12, or Adpgk peptides, respectively. Each condition has controls without peptide and without cancer cells (+ or - peptide). N=6 for each condition and the mean and SD from the mean are shown.

### 3 out of the 4 new MC38 TCRs are H2-K^b^-dependent

Experiments with the previously identified MC38 neoantigens suggested that the Adpgk-derived peptide may be the cognate antigen for one out of the four MC38 TCRs (TCR 30). In an effort to de-orphanize the other three TCRs, we conducted exome sequencing of the MC38 cell line and predicted neoantigens using the MHCFlurry pipeline (see Methods for details). Our goal was to screen the MC38 TCRs with peptides for the first 100 neoepitopes with highest predicted affinity. To narrow down the list of peptides to screen, we determined which allele the MC38 TCRs were dependent on for peptide presentation (H2-K^b^ or H2-D^b^). To achieve this we used CRISPR/Cas9 to knock out H2-K^b^ or H2-D^b^ from MC38 cells and achieved about 95% efficiency for each allele (shown in Figure 8A and B). We co-cultured Jurkat-NFAT cells expressing the MC38 TCRs with the MHC allele knockout MC38 cells and measured T cell activation via luciferase luminescence. We included OT-I TCR expressing cells as a control, with or without SIINFEKL peptide addition, because SIINFEKL is dependent on H2-K^b^ presentation. We observed the expected decrease in activation of the OT-I-expressing cells with SIINFEKL when co-cultured with the H2-K^b^ deficient MC38 cells (Figure 8C, pink bar). Likewise, our results indicated that MC38 TCRs 31, 32, and 33 are H2-K^b^ dependent. The results were not conclusive for TCR30, but because we had already identified the cognate antigen for this TCR (which is predicted to be presented by H2-D^b^), we limited our list of neoantigen peptides to screen to those predicted to be presented by the H2-K^b^ allele. The full list of 96 peptides we screened is provided as a supplemental table.

**Figure 8.**
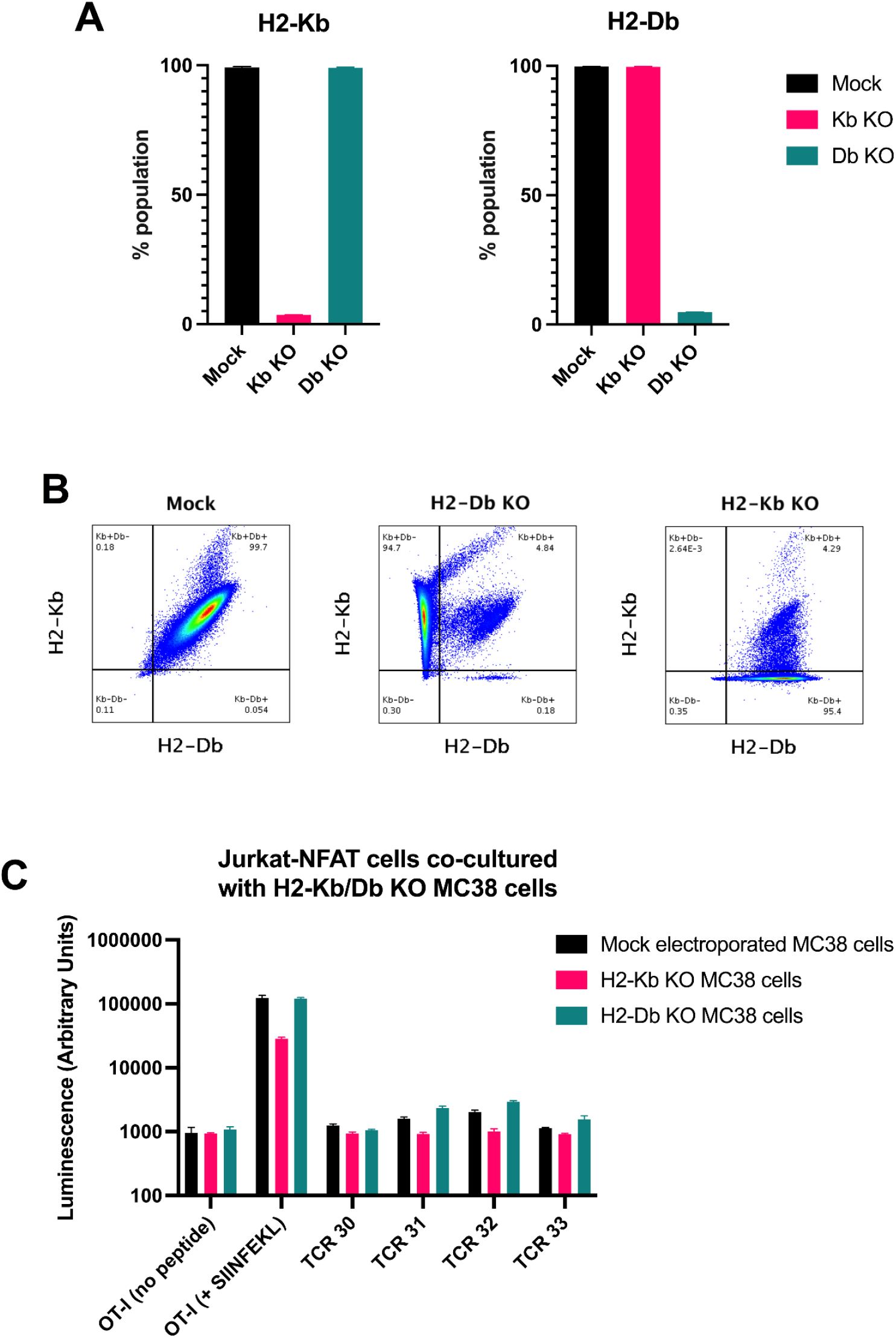
MC38 TCRs 31, 32, and 33 appear to be dependent on H2-K^b^ antigen presentation. A) MC38 cells were electroporated with sgRNAs and Cas9RNP against H2-K^b^ (pink bars), H2-D^b^ (teal bars), or mock conditions (black bars) and assayed for KO efficiency at day 3 post-electroporation. Percentage of live population positive for H2-K^b^ (left graph) and H2-D^b^ (right graph) is shown for each condition. B) Representative flow cytometry plots of each condition. H2-D^b^ staining on the x-axis and H2-K^b^ staining on the y-axis. C) Jurkat-NFAT cells expressing either the OT-I TCR or one of the MC38 TCRs were co-cultured with MC38 cells for 12 hours before being assayed for luciferase expression on a luminescence-capable plate reader. The positive control is OT-I pulsed (indicating the addition of the SIINFEKL peptide to the cultures with the Jurkat-NFAT cells expressing the OT-I TCR) and the negative control is OT-I unpulsed, indicating no addition of SIINFEKL peptide (black bars on A-D). Pink bars indicate conditions with one of the MC38 TCRs. The dotted black line is drawn at the average activation for each TCR co-cultured with the MC38 cells alone (i.e. no peptide was added). N=6 for each condition and the mean and SD from the mean are shown.

### MC38 TCRs 31-33 remain orphan TCRs

The results of the first peptide screen with the T cell activation bioassay are shown in Figures 9 and 10. Each TCR was on a separate plate with the 96 peptides, and there were 3 replicate plates per TCR. Controls were on a separate plate. We included the Adpgk-derived peptide in our screen to verify our previous results shown in Figure 7. We observed the same increase with the Adpgk-derived peptide in the cultures with TCR 30, but not in any of the cultures with the other 3 Neo TCRs. The vertical pink dotted lines in the graphs in Figures 9 and 10 indicate the luminescence level of the MC38 TCR co-cultured with the MC38 cells alone. Again, we were looking for any peptides that resulted in increased luminescence with any of the TCRs.

**Figure 9.**
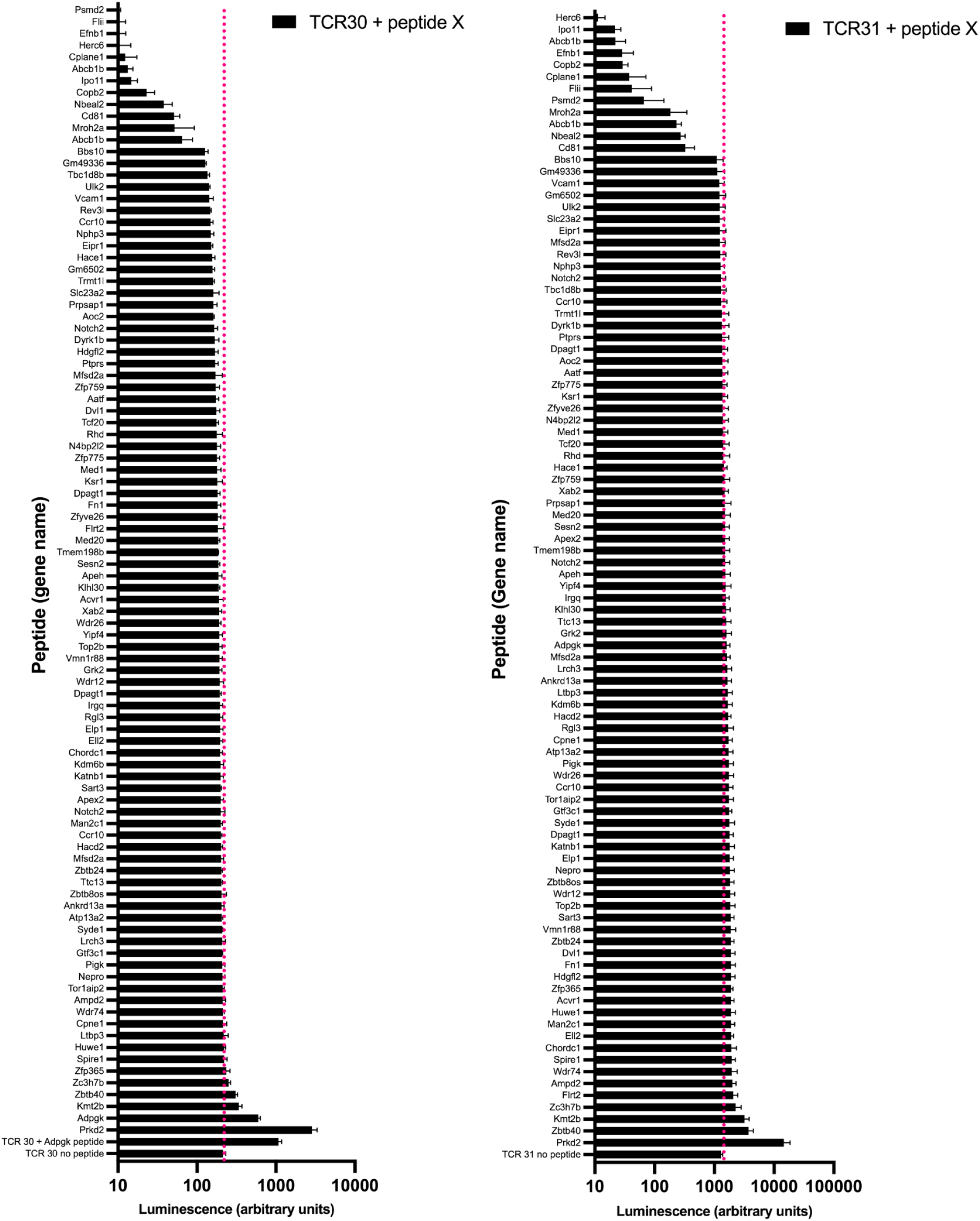
T cell activation bioassay with MC38 TCR 30 and TCR 31 in co-culture with 96 predicted MC38 neoantigen peptides. The final peptide concentration used was approximately 200 uM and the co-culture incubated for 12 hours before luciferase expression was measured via luminescence on a plate reader. n=3 for each condition. White opaque plates with clear bottoms were used for all TCR 30 conditions (including controls). White opaque plates with opaque bottoms were used for TCR 31 conditions (including the OT-I TCR controls). The use of different plates for the TCR30 samples was due to COVID-19-related plate shortages.

**Figure 10.**
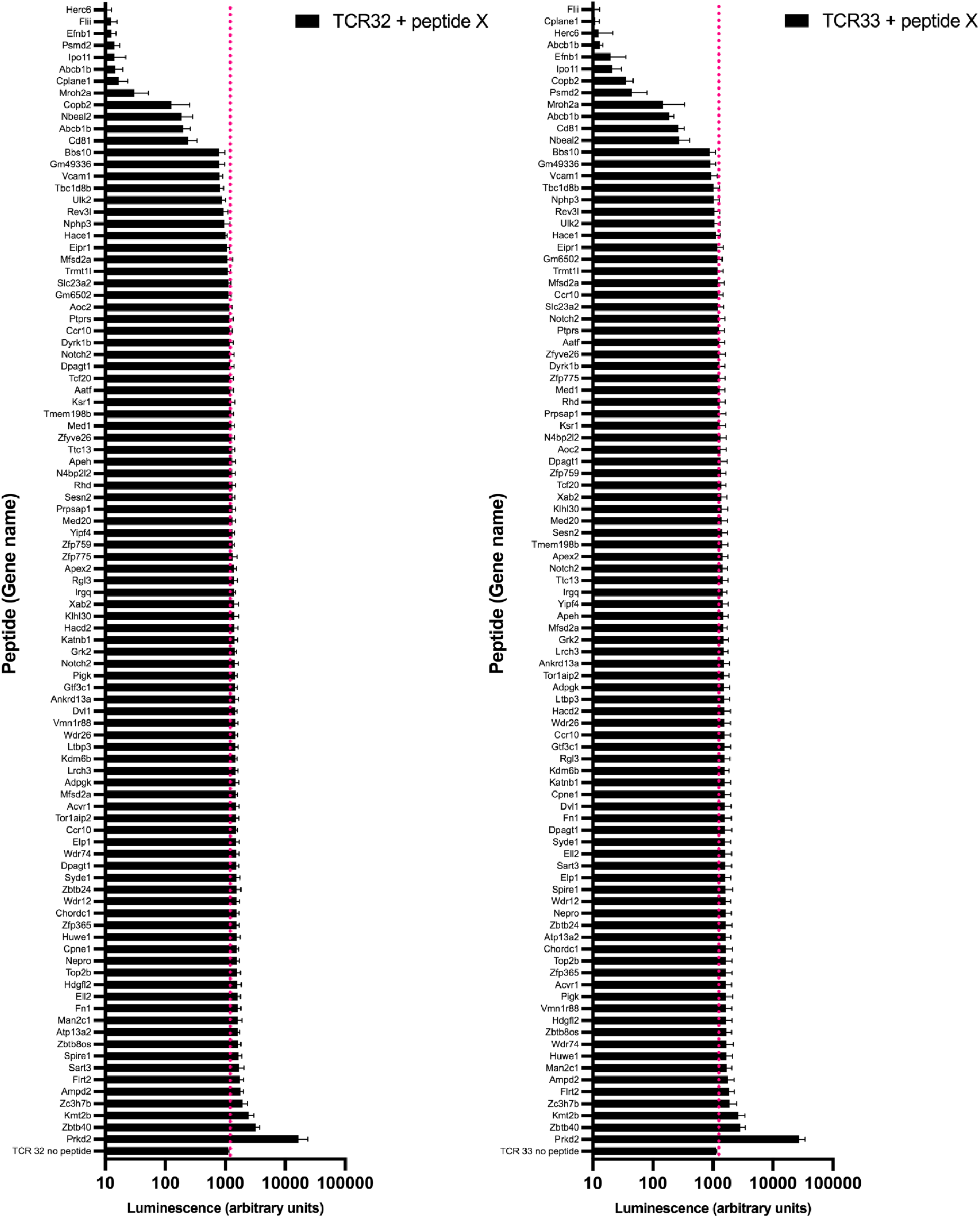
T cell activation bioassay with MC38 TCR 32 and TCR 33 in co-culture with 96 predicted MC38 neoantigen peptides. The final peptide concentration used was approximately 200 uM and the co-culture incubated for 12 hours before luciferase expression was measured via luminescence on a plate reader. n=3 for each condition. White opaque plates with opaque bottoms were used for TCR 32 and TCR 33 conditions (including the OT-I TCR controls).

For any peptides that tested positive, we repeated the assay with more replicates. The results for the repeated experiments are shown in Figure 11. There was one peptide in particular (derived from the Prkd2 gene) that resulted in an increase with all MC38 TCRs in the first screen, which led us to include extra controls by culturing OT-I and Pmel TCR-expressing cells with the neoantigen peptides (Figure 11, bottom two graphs). The positive control was the SIINFEKL peptide for the OT-I TCR and the hgp100 peptide for the Pmel TCR. Again, the Adpgk-derived peptide only activated TCR 30-expressing Jurkat cells. The Prkd2-derived peptide activated all TCRs, including the OT-I and Pmel TCRs. None of the remaining peptides activated MC38 TCRs 31-33.

**Figure 11.**
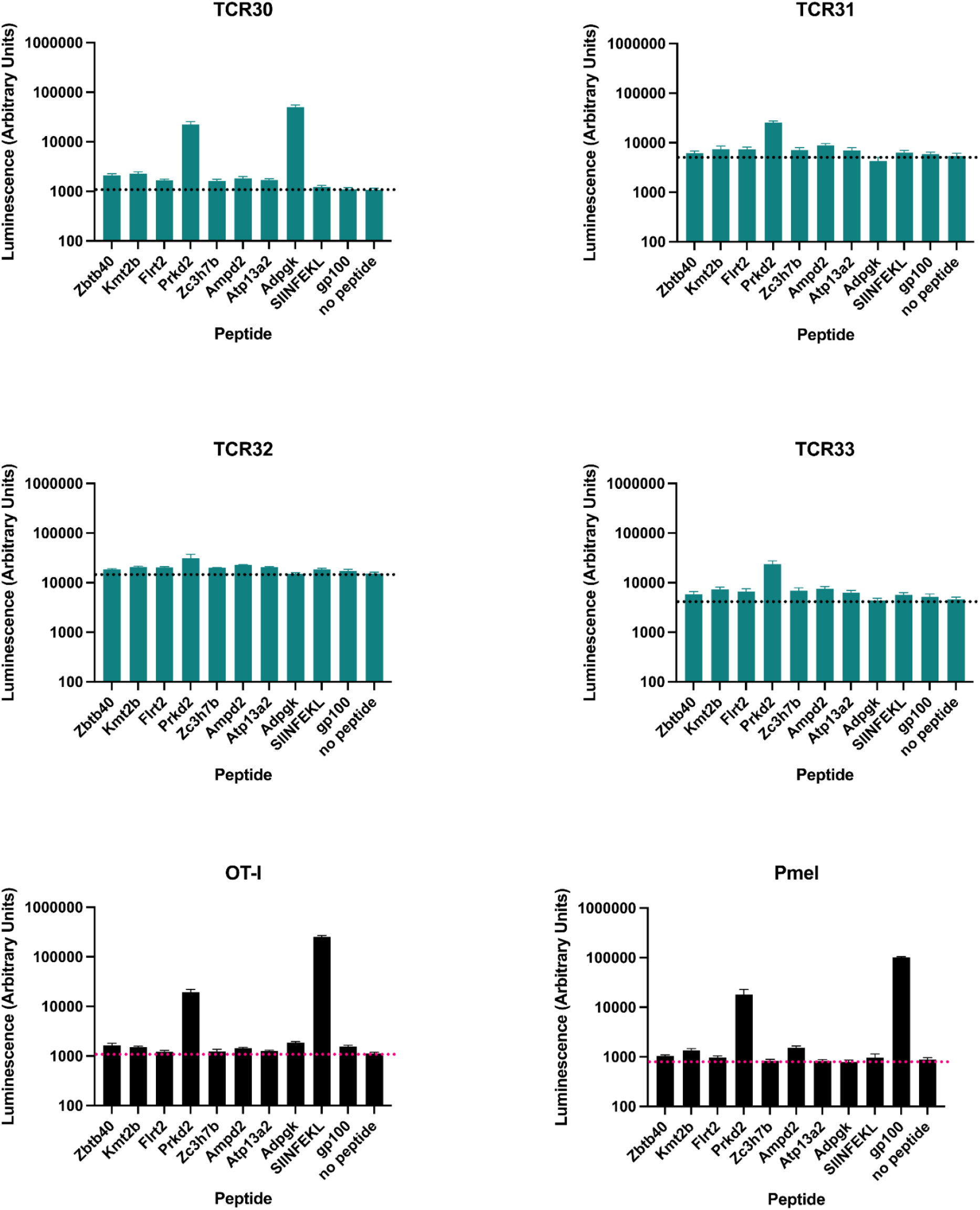
T cell activation bioassay with MC38 TCRs and possible activating peptides. Peptides that showed an increase in luminescence in the first screen (Figures 9 and 10) were repeated in co-culture with MC38 cells and the MC38 TCRs or the OT-I or Pmel-1 TCRs. Final peptide concentration was approximately 100 uM. The co-culture ran for 12 hours before the luciferase expression was measured via luminescence. White bottomed opaque 96 well plates were used for all samples. n=4 for each condition.

In the screen with the 96 peptides, we observed that several peptides resulted in markedly reduced levels of luminescence (in Figures 9 and 10). We thought it may be due to the solvent having a detrimental effect on the cells. We repeated the experiments with these peptides, diluting them 1:10 in culture media (results shown in Figure 12). None of these peptides resulted in an increase in luminescence with any of the MC38 TCRs.

**Figure 12.**
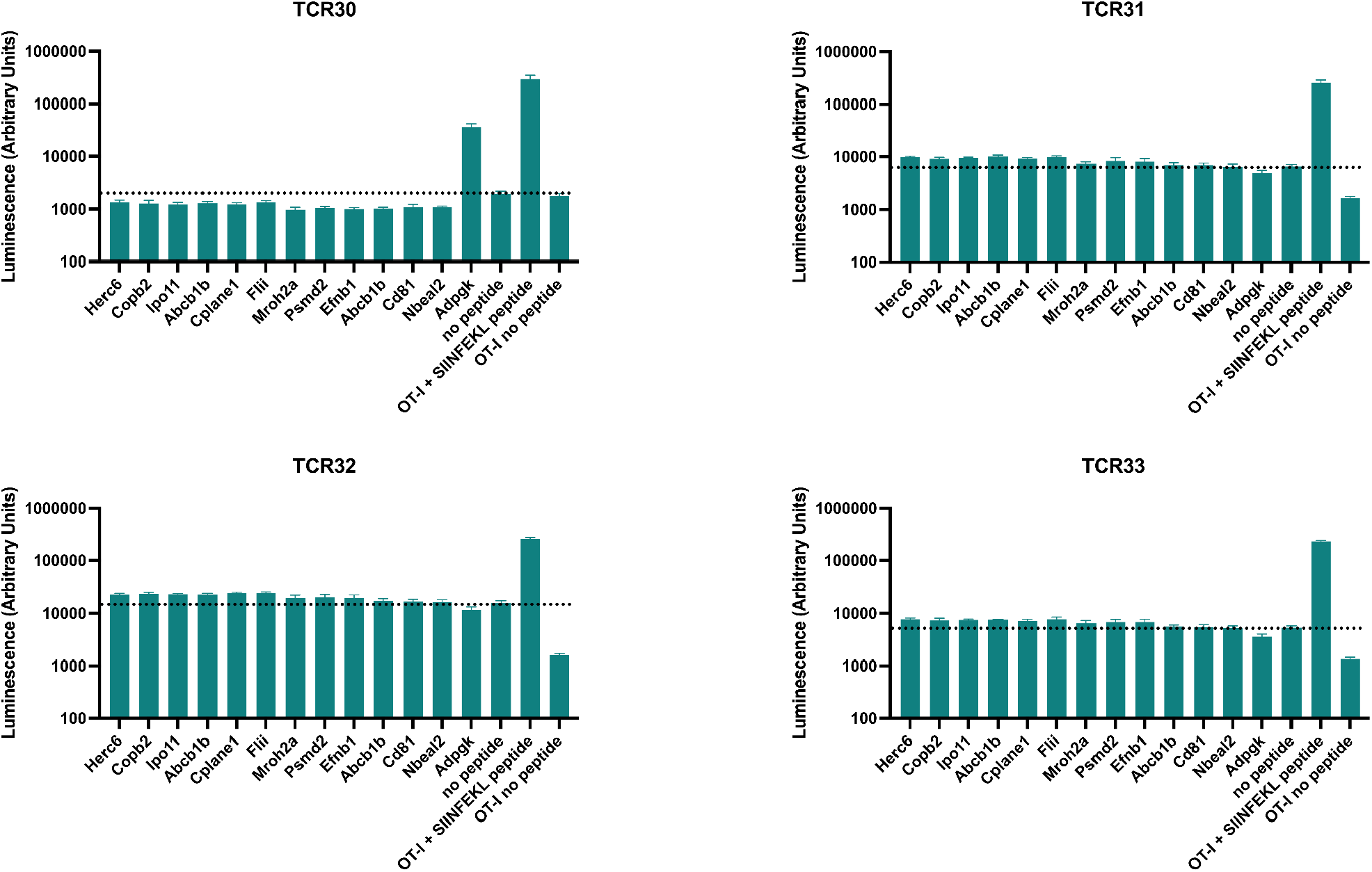
T cell activation bioassay with MC38 TCRs and peptides from the full screen in Figures 9 and 10 that resulted in low signal. Peptides from the full screen (shown in Figures 9 and 10) that resulted in a decrease in luminescence compared to the “no peptide” condition were diluted 1:10 in cell culture media before co-culturing with MC38 cells and MC38 TCR-expressing Jurkat-NFAT cells. The co-culture ran for 12 hours before the luciferase expression was measured via luminescence. The OT-I TCR was used a positive and negative control with and without the addition of the SIINFEKL peptide, respectively. White bottomed opaque 96 well plates were used for all conditions. n=4 for each condition.

### LLC-A9F1 tumor TIL-derived TCRs fail to activate Jurkat-NFAT cells when co-cultured with LLC-A9F1 cells

One of our primary intentions in this investigation was to show that our approach to TCR discovery and reactivity testing would extend to a second tumor model. We started with two cell lines (MC38 and LLC-A9F1) and conducted the tumor growth, TIL isolation, and immune profiling simultaneously. Once we had the TCR constructs, we began functional experiments with the MC38 TCRs simply because the MC38 cell line is easier to handle than the LLC-A9F1 cells. MC38 cells are purely adherent, whereas as LLC-A9F1 are a mixed population of adherent and suspension cells. Additionally, cultured LLC-A9F1 cells are known to have low MHC class I expression, resulting in the cells being poor targets for T cells (Eisenbach, Segal, and Feldman 1983). Treatment with IFNγ is used to increase the cell surface levels of MHC class I, which can allow for greater T cell targeting. We tested out IFNγ treatment in LLC-A9F1 cells and observed a significant increase in H2-K^b^ and H2-D^b^ surface expression with overnight treatment and a range of concentrations of IFNγ (data not shown). We used the T cell bioactivity assay to test the LLC-A9F1 TCRs (results shown in Figure 13). We included IFNγ in the culture media (at a concentration of 10 ng/ml) when seeding the LLC cells the day before the co-culture with the LLC TCR-expressing T cells. Because we were not sure of the optimal time for the T cells to be activated by the LLC-A9F1 cells, we allowed the co-cultures to proceed for 5 hours and for 12 hours before measuring luminescence. The OT-I TCR with and without the addition of SIINFEKL served as the positive and negative controls, respectively. As shown in Figure 13, cells expressing the OT-I TCR were activated when the SIINFEKL peptide was added to the co-culture with the LLC-A9F1 cells after both 5 hours and 12 hours duration, but not when the SIINFEKL peptide was omitted. However, none of the LLC-A9F1 TCRs resulted in T cell activation.

**Figure 13.**
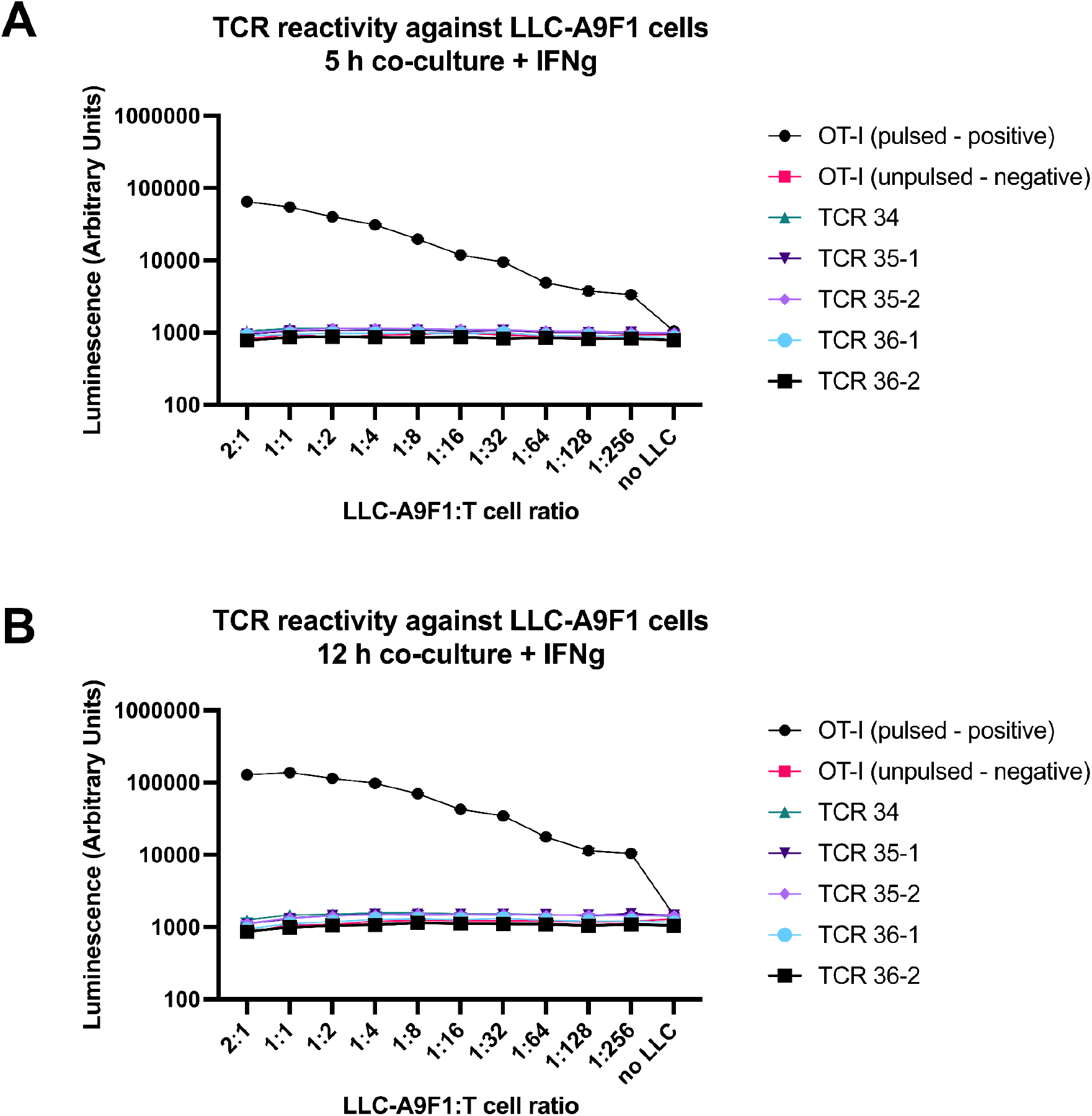
LLC-A9F1 TCRs do not result in T cell activation when co-cultured with LLC-A9F1 cells. Jurkat-NFAT cells expressing either the OT-I TCR or one of the LLC-A9F1 TCRs were co-cultured with IFNγ-treated LLC-A9F1 cells for 5 (A) or 12 (B) hours before luciferase expression was measured on a luminescence-based plate reader. The positive control is OT-I pulsed (indicating the addition of the SIINFEKL peptide to the cultures with the Jurkat-NFAT cells expressing the OT-I TCR) and the negative control is OT-I unpulsed, indicating no addition of SIINFEKL peptide (black and pink lines, respectively). N=6 for each condition and the mean and SD from the mean are shown.

In the experiments in Figure 13, we aspirated the culture media from the wells of LLC-A9F1 cells on the day of co-culture in order to add the T cell suspension in fresh culture media. This would have likely removed LLC-A9F1 cells that were growing in suspension. We repeated the experiment with 12 hour co-culture, but this time we centrifuged the plates of LLC-A9F1 cells before aspirating the culture media and adding the T cells, to ensure that both populations of LLC-A9F1 cells were included in the co-culture. Results in Figure 14 show that there was still no T cell activation with any of the LLC-A9F1 TCRs with this further measure.

**Figure 14.**
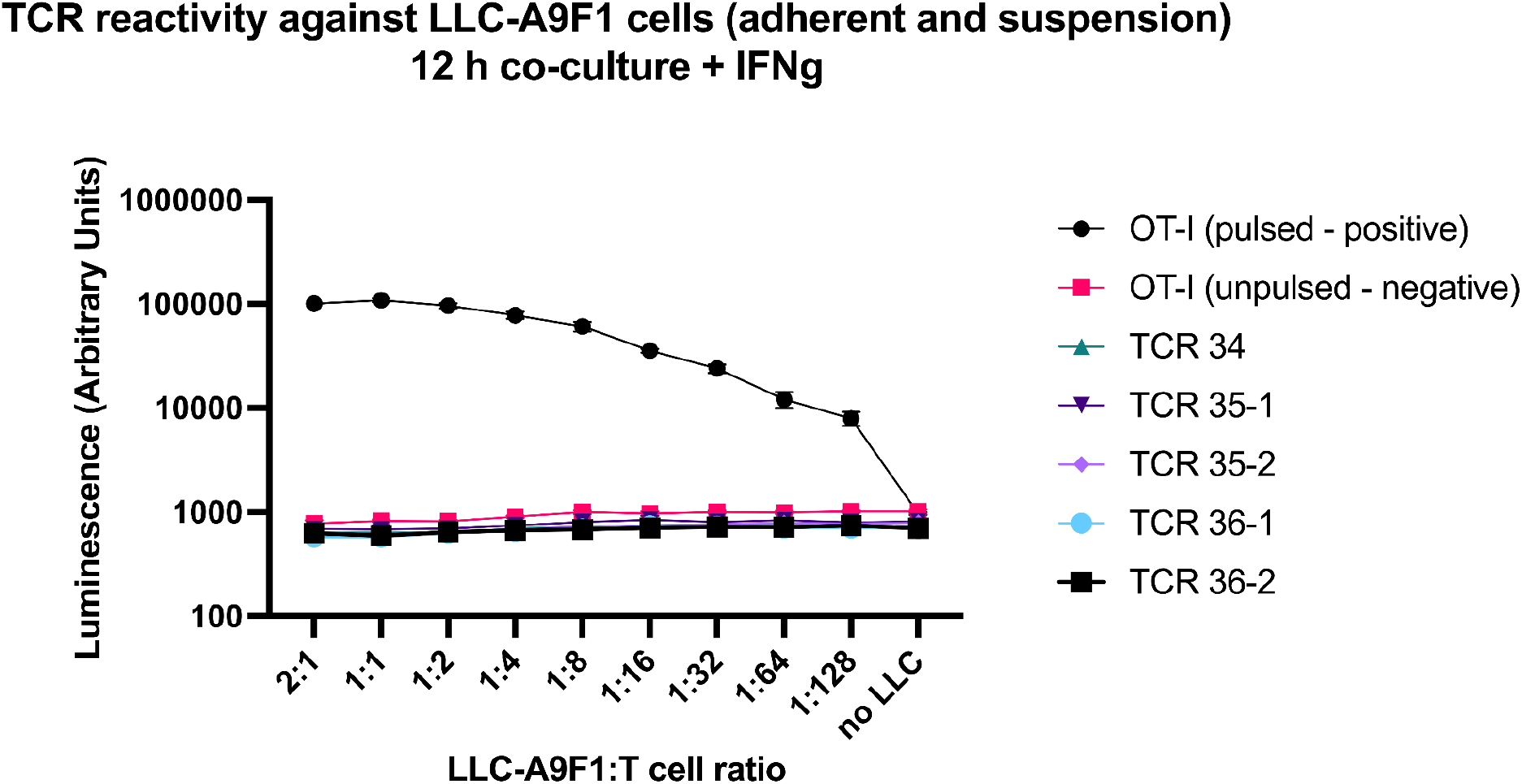
LLC-A9F1 TCRs do not result in T cell activation when co-cultured with LLC-A9F1 cells (including adherent and suspension cells). Jurkat-NFAT cells expressing either the OT-I TCR or one of the LLC-A9F1 TCRs were co-cultured with IFNγ-treated LLC-A9F1 cells for 12 hours. The plates with LLC-A9F1 cells were centrifuged before aspirating culture media and adding the T cells. Luciferase expression was measured on a luminescence-based plate reader. The positive control is OT-I pulsed (indicating the addition of the SIINFEKL peptide to the cultures with the Jurkat-NFAT cells expressing the OT-I TCR) and the negative control is OT-I unpulsed, indicating no addition of SIINFEKL peptide (black and pink lines, respectively). N=6 for each condition and the mean and SD from the mean are shown.

## Discussion

In this work we have shown that it is possible to obtain paired tumor-reactive TCR sequences directly from freshly isolated TILs via single cell immune profiling and to quickly test the reactivity of the newly identified TCRs by expressing them in other T cells using mRNA electroporation and co-culture systems. In the case of the MC38 TCRs, we have shown that when expressed in T cells, they have the ability to recognize and eliminate parent tumor cells and to mediate T cell activation. Additionally, we have paired a previously predicted neoantigen (ASMTNMELM derived from the Adpgk gene) with one of the MC38 TCRs, TCR30 (or Picky Sticky). TCR30 may be useful for the development of an MC38 TCR transgenic mouse model.

We made some additional observations through the course of our investigation. The single cell immune profiling results for the MC38 samples showed that each tumor resulted in a different dominant TCR clonotype, even within the consistency of a mouse model. But, the most frequent TCR isolated from one tumor was found at a low frequency in other samples, which indicates that the most frequent TCR pair is not necessarily the most reactive and that there may be many different tumor-reactive clones at varying abundances in a TIL population. These observations could suggest that TIL expansion in order to obtain sufficient numbers of cells for therapy may alter the composition of the TCRs in the starting TIL population, potentially resulting in the loss of more tumor-reactive clones. In one study, when multiple independent TIL cultures were initiated from a single tumor, each culture resulted in a different dominant TCR and each dominant TCR exhibited a different level of reactivity to tumor cell lines (Dudley et al. 2003). In that study it was found that each independent TIL culture had a different reactivity to the tumor lines they were tested against, suggesting different clones had expanded in each culture. Furthermore, when several populations of TILs were expanded independently from a single excisional biopsy and then combined for cell therapy, the resulting cell product had increased tumor-reactivity compared to when a single culture from a tumor was expanded. Our results align with these observations at the TCR sequence level. The approach we took could potentially be used as a tool to study the evolution of TCR clonotype frequency in *ex vivo* TIL expansion. Finally, our experiments revealed that one of the peptides from our MC38 neoantigen prediction (derived from the Prkd2 gene) resulted in T cell activation with any TCR that we expressed in the Jurkat-NFAT cells, including the OT-I and Pmel TCRs, when co-cultured with MC38 cells. Further work is needed to determine if this peptide is an indiscriminate activator of T cells and the mechanism behind the activation.

We recognize there are limitations to our system. First, we did not knock out the endogenous TCR in the human donor T cells or the Jurkat-NFAT cells before expressing the exogenous TCRs. There is evidence that endogenous TCR expression can reduce the surface expression of the desired introduced TCR due to heterodimerization with endogenous subunits and competition for CD3 (Heemskerk et al. 2007). However, there is also evidence that when a TCR with a murine constant region is expressed in human T cells (such as with our set-up), that the murine TCR is more efficient at interacting with CD3 than the endogenous human TCR, resulting in increased murine TCR surface expression (Cohen et al. 2006). In any case, even with potential interference from endogenous TCR expression in our system, we were able to detect a response both in our cytotoxicity assays and T cell activation bioassays with the MC38 TCRs. Since we were not successful in detecting a response with the LLC-A9F1 TCRs, modifying the system by knocking out endogenous TCRs is an approach to consider in the future. Secondly, we used MC38 cells as the antigen-presenting cells in our experiments to test the MC38 TCRs because this is what we had on hand. It was necessary to initially show that the TCRs were reactive against the cell lines that were used for their parental tumors. While this approach was informative for our purposes, this is likely not an ideal set-up for TCR deorphanization by co-culture with predicted neoantigen peptides because there is less of a difference in signal between the “no peptide” condition and any condition with a peptide. Consequently, a signal from a weak TCR-peptide interaction may not be detected. Adapting our system to use a different target cell line may be beneficial, especially for further work with the three remaining orphan MC38 TCRs and the LLC-A9F1 TCRs.

## Material and Methods

### Cells, cell lines, and animal model

#### Human primary T cells

Buffy coat from healthy human donors was purchased from Plasma Consultants LLC, Monroe Township, NJ. T cells were isolated with the EasySep™ Direct Human T Cell Isolation Kit (StemCell, Vancouver, Canada). Isolated T cells were cultured in media containing: RPMI with L-glutamine (Corning), 10% fetal bovine serum (Atlas Biologicals, Fort Collins, CO), 715 uM 2-mercaptoethanol (EMD Millipore), 25 mM HEPES (HyClone, GE Healthcare, Chicago, IL), 1% Penicillin-Streptomycin (Gibco), 1X sodium pyruvate (HyClone, GE Healthcare, Chicago, IL), and 1X non-essential amino acids (HyClone, GE Healthcare).

#### Cell lines

MC38 mouse colon adenocarcinoma cells and LLC-A9F1 (Lewis Lung Carcinoma - clone A9F1) cells were kindly provided by the Rubinstein and Wrangle lab at the Medical University of South Carolina. The MC38 cells were kindly provided to the Rubinstein and Wrangle lab by Aaron Ring and were originally from the Marcus Bosenberg Lab. Jurkat-NFAT cells were purchased from Promega (Madison, WI) as part of the T Cell Activation Bioassay kit. MC38, LLC-A9F1, and Jurkat-NFAT cells were cultured in RPMI 1640 with L-Glutamine (Corning, Durham, NC) supplemented with 10% Fetal Bovine Serum (Atlas Biologicals, Fort Collins, CO) and 1% Penicillin-Streptomycin (Gibco).

#### Mice and tumor protocol

Female C57BL/6 (H2-K^b^/H2-D^b^) mice were used as the host for the MC38 or LLC-A9F1 tumors. 300,000 MC38 cells or 1 million LLC-A9F1 were subcutaneously injected into 16 mice (8 mice per cell line). Tumors were monitored every 1 to 3 days and allowed to grow until they reached the size of 100 mm^3^, after which the tumors were excised and directly processed for TIL isolation. Tumor growth curves can be viewed here: https://github.com/hammerlab/tcr-transfer/blob/master/analyses/Explore%20tumor%20growth%20curves%20for%20MC38%20and%20LLC%20cell%20lines.ipynb

### CD8+ TIL isolation from tumor samples

Each tumor was dissociated into a single cell suspension with Miltenyi’s gentleMACS Dissociator according to the tumor dissociation kit protocol (Miltenyi Biotec, Germany). A small portion of the dissociated tumor cells was removed for profiling via flow cytometry. The remaining cell suspension was used to isolate CD8+ T cells with the magnetic-bead-based positive selection kit: Dynabead’s positive CD8+ isolation kit (Thermo Fisher, Waltham, MA). It is important to use an isolation kit that allows separation of cells from the beads for the samples to be compatible with 10x’s single-cell preparation. The isolated CD8+ cells were kept in T cell media supplemented with IL-2 at 200 IU/ml (see culture media components in the primary human T cell methods section) until the samples were prepared for scTCR sequencing (which happened on the same day as tumor dissociation and CD8+ isolation). To determine which samples to process for sequencing, the isolated CD8+ cells were inspected visually with a light microscope and cell count and viability was estimated with Trypan Blue staining. We prioritized samples that had >55-60% viability and ones where the average cell size was 10-11 microns. From a total of 8 tumors for each cell line, we chose the 4 with the highest viability from each cell line and continued with the TCR library preparation on the same day of the isolation.

### Emulsion and TCR library preparation for single-cell sequencing

Experimental procedures followed established techniques using the Chromium Single Cell 5’ Library & Gel Bead Kit (10x Genomics; CG000086 Rev L). Briefly, magnetically enriched CD8+ T cells in single-cell suspension were loaded onto a 10x Genomics Chip A and emulsified with 5’ Single Cell GEM beads using a Chromium™ Controller (10x Genomics). TCR libraries were constructed from the barcoded cDNAs using the 10x Genomics Chromium™ Single Cell V(D)J Enrichment Kit, Mouse T Cell (Translational Science Laboratory at the Medical University of South Carolina). Sequencing was performed on each sample (approximately 50 million reads/sample) using a NovaSeq S4 flow cell (Illumina) at the VANTAGE facility (Vanderbilt University Medical Center).

### Plasmids

#### TCR plasmid construct design

The 10x V(D)J sequencing results showed TCR enrichments based on the unique CDR3 sequence for each alpha and beta chain. Therefore, it was necessary to complete the TCR sequence by adding the constant region before cloning it into a mammalian expression vector and ultimately to be able to express a functional TCR. To construct the full TCR for cloning, the TCR sequencing results were visualized in the Loupe V(D)J Browser (free software from 10x Genomics, Pleasanton, CA) to find the most abundant clonotype in each tumor. The consensus sequences for the unique CDR3 in each alpha and beta chain from the clonotype of interest were exported in FASTA file format. The coding region was identified in the consensus sequence from the start codon ATG. The 10x VDJ sequencing often does not include the whole constant region. Therefore, the sequences for the alpha and beta constant regions from the background of the mouse used to bear the tumors (C57BL/6) were used to manually complete the missing 3’ portion of the TCR sequence. Gene synthesis and cloning was performed by GenScript (Piscataway, NJ).

Step-by-step extraction of the corresponding TCR chain sequence from the single-cell sequencing results and cloning them into the vector is available at https:/ob/master/other/HANDLING_TCR_SEQUENCES.md. In short, we extracted the consensus alpha and beta chain sequences of the most abundant clones using the Loupe browser; and then completed the constant chain sequence based on the TRAC and TRBC reference sequences. Once we had the complete TRA and TRB sequences constructed, we computationally identified cutting sites that are compatible with the pcDNA3.1 vector and that would not cut any of the candidate TCR sequences: https://github.com/hammerlab/tcr-transfer/blob/master/analyses/Pick%20the%20best%20enzymes%20for%20cloning%20all%20TCRs%20into%20the%20plasmid.ipynb. Based on this analysis we picked AflII and NotI for the cloning sites as none of the TCR sequences had the cutting site for both of these enzymes, providing a uniform cloning strategy across all TCR sequences.

#### Plasmids

DNA plasmids were transformed in Max Efficiency DH5α Competent cells (Thermo Fisher Scientific, Waltham, MA), propagated in Luria Broth, and purified using Qiagen midi-prep kits (Hilden, Germany).

The mouse CD8a and CD8b plasmids were purchased from Origene (Rockville, MD). Catalog numbers MR227539 and MR225204, respectively.

We deposited the following plasmids to Addgene:

- pcDNA3.1(+) OTI-TCRA (Plasmid #131035)
- pcDNA3.1(+) OTI-TCRB (Plasmid #131036)
- pcDNA3.1(+) Pmel-1-TCRA (Plasmid #175287)
- pcDNA3.1(+) Pmel-1-TCRB (Plasmid #175288)
- pcDNA3.1(+)-MC38-TCRA-30-picky_sticky (Plasmid #138983)
- pcDNA3.1(+)-MC38-TCRB-30-picky_sticky (Plasmid #138984)
- pcDNA3.1(+)-MC38-TCRA-31-murderous_crow (Plasmid #138985)
- pcDNA3.1(+)-MC38-TCRB-31-murderous_crow (Plasmid #138986)
- pcDNA3.1(+)-MC38-TCRA-32-riot_punisher (Plasmid #138987)
- pcDNA3.1(+)-MC38-TCRB-32-riot_punisher (Plasmid #138988)
- pcDNA3.1(+)-MC38-TCRA-33-MC_hammer (Plasmid #138989)
- pcDNA3.1(+)-MC38-TCRB-33-MC_hammer (Plasmid #138990)

### In vitro transcription (IVT)

Plasmids were linearized overnight with restriction enzymes (NotI for mCD8 and TCR constructs, EcoRI for OT-I constructs) (New England BioLab, Ipswich, MA). The linearized DNA was cleaned up via phenol chloroform extraction and sodium acetate and ethanol precipitation prior to IVT. mRNA was generated from the T7 promoter in the linearized plasmids with the HiScribe T7 ARCA mRNA kit with tailing (NEB #E2060S, Ipswich, MA). We would typically use half of a kit (10 reactions) per TCR subunit with 10 ug of template DNA, which would result in approximately 300 - 450 ug of mRNA resuspended in RNA elution buffer (0.1 mM EDTA) with 10 minutes incubation at 65°C. RNA concentration was measured on the NanoDrop One.

### TCR electroporation

Human primary T cells were isolated from healthy human donor buffy coats and cultured as described above. Directly after isolation, T cells were activated with anti-CD3/CD28 magnetic dynabeads (Thermo Fisher) at a bead to cell concentration of 1:1, with a supplement of 200 IU/ml of IL-2 (NCI preclinical repository). Cells were debeaded after two days of activation using DynaMag (Thermo Fisher) magnetic tube holders to remove the beads. The remaining cells were enriched for CD8+ cells with Dynabeads™ Untouched™ Human CD8 T Cells Kit (Thermo Fisher #11348D) following the manufacturer’s protocol. Initially, we attempted to directly isolate CD8+ T cells from the buffy coat but found the viability of the cells was better when activated as pan T cells and then enriched for CD8+ cells.

Directly after CD8 enrichment, the T cells were electroporated. The cells were centrifuged for 5 minutes at 350 x g and washed 3 times with PBS before being resuspended in R buffer (a component of the Neon electroporation kit, Thermo Fisher). Cells were resuspended in R buffer at a concentration of 20 million cells per ml, taking into account the volume of the RNA mix to be added later. The Neon 100 ul tip was used, which meant 2 million cells were electroporated per reaction with 10 ug total of RNA (5 ug of each TCR subunit). The electroporation settings used were: voltage of 1600, 10 ms width, and 3 pulses.

Jurkat-NFAT cells were centrifuged for 5 minutes at 350 x g and washed 3 times with PBS before being resuspended in R buffer. The Neon 100 ul tip was used to electroporate 2 million cells per reaction with 20 ug total RNA (5 ug each TCR subunit and 5 ug each mCD8 subunit) Jurkat-NFAT cells were electroporated at a voltage of 1350 with a pulse width of 10 and 3 pulses.

### Flow Cytometry

Flow Cytometry was performed on the BD FACSVerse or LSR Fortessa Flow Cytometers. Cells were harvested, centrifuged for 5 minutes at 300 x g, and resuspended in PBS + 20% FBS. Antibodies were added at the recommended concentrations and stained at room temperature for 15 minutes, protected from light.

#### Multimer staining for flow cytometry

Jurkat-NFAT cells expressing the OT-I TCR via mRNA electroporation were stained with flow buffer containing the OT-I dextramer (APC) for 15 minutes before adding antibodies against mCD8A (Pe-Cy7) and mCD8B (V421), and then stained for an additional 15 minutes.

### OT-I Dextramer

| Name | Vendor | Catalog number |
| --- | --- | --- |
| H2-K<sup>b</sup> SIINFEKL OVA dextramer | Immudex | JD2163 |

### Antibodies

| Antibody name | Vendor | Catalog number |
| --- | --- | --- |
| Anti-mouse H2-K<sup>b</sup> (APC) | Biolegend | 116518 |
| Anti-mouse H2-D<sup>b</sup> (PerCy5.5) | Biolegend | 111518 |
| Anti-mouse CD8a (Pe-Cy7) | Biolegend | 100722 |
| Anti-mouse CD8b (V421) | Biolegend | 126629 |

### Flow cytometry-based cytotoxicity assay

Full protocol is available at: dx.doi.org/10.17504/protocols.io.83bhyin

### T Cell Activation Bioassay

The T Cell Activation Bioassay kit was purchased from Promega. The Jurkat-NFAT cells were thawed and cultured in RPMI (supplemented with 10% FBS and 5% pen/strep) as per the kit protocol. After the initial thawing, we expanded the cells and froze vials for further experiments.

On day 1 of the experimental setup, MC38 or LLC-A9F1 cells were plated either at 100,000 cells per well, or a 1:2 serial dilution was performed starting from 200,000 cells per well. The target cells were plated in opaque, white 96 well plates for luminescence (Griener BioSciences, Monroe, NC). On the same day, Jurkat-NFAT cells were electroporated with mRNA encoding the TCRs of interest and mCD8 subunits. The electroporated Jurkat cells were cultured overnight in 6 well plates or T75 flasks in RPMI supplemented with FBS (the pen/strep was omitted).

The next day (at least 12 hours later, usually closer to 24 hours) the Jurkat-NFAT cells were counted, centrifuged, and resuspended in fresh culture media at a concentration to achieve 100,000 cell per well. The media in the plates with the target cells was aspirated and the Jurkat cell suspension was added to each well. The cells were co-cultured for 6 or 12 hours before adding the T Cell Activation Bioassay substrate/assay buffer. Once the buffer was added the plates were incubated at room temperature, protected from light, for 10 minutes before the luminescence was measured with the SpectraMax iD5 plate reader (Molecular Devices, San Jose, CA).

The full protocol is available at: dx.doi.org/10.17504/protocols.io.ba8gihtw

### MC38 RNA and whole exome sequencing and neoepitope prediction

MC38 cells and PBMCs from a C57BL/6 mouse (to use as the normal sample) were pelleted and submitted to GeneWiz (South Plainfield, NJ) for RNA- and WEX-sequencing. Mutations were called using OpenVax’s Neoantigen Vaccine Pipeline (https://github.com/openvax/neoantigen-vaccine-pipeline), where mouse PBMC WEX-Seq data was used as a normal and the MC38 data as the matched tumor sample based on the mm10 reference genomes. Once the mutations were called, Strelka and MuTeCT outputs were then fed to Topiary (https://github.com/openvax/topiary) using MHCflurry (O’Donnell, Rubinsteyn, and Laserson 2020) as the back-end for neoantigen prediction (https://github.com/openvax/mhcflurry). RNA-seq data was quantified and summarized on the gene level using kallisto (https://github.com/pachterlab/kallisto/) based on Ensembl’s mouse cDNA reference sequence (v103 - GRCm39). Raw, intermediate, and merged data files are available at https://github.com/hammerlab/tcr-transfer/tree/master/analyses/neoantigen/mc38b. The neoantigen peptide list was limited to peptides the length of 9 amino acids, with predicted H2-K^b^-dependency, and with transcripts per million (TPM) greater than 1. The resulting list was ranked by affinity, and the top 96 peptides were chosen for the screen. The list is provided in supplemental data.

### Peptides

Peptides were purchased from Genscript. Previously predicted peptide sequences for MC38 neoantigens were used from the cited publications (Yadav et al. 2014; Hos et al. 2019). Further neoantigens were predicted for the MC38 cell line we used in our experiments using MHCFlurry as described above. Peptides were reconstituted in solvents recommended by Genscript through the peptide solubility test service, aliquoted, and stored at −80. For the large peptide screen 0.5 mg of each peptide was provided in a 96-well format. The peptides were reconstituted in 50 ul of recommended solvent and pipetted directly into the co-cultures in 96 well plates. The remaining peptide was stored at −80. A complete list of peptides and solvent used for each is provided in supplemental data.

### H2-K^b^ and H2-D^b^ knockout in MC38 cells

The sgRNAs and Cas9RNP were purchased from Synthego (Menlo Park, CA). We used the Synthego CRISPR Design Tool to design the sgRNA sequences. We resuspended the sgRNAs in TE buffer before use. The sgRNAs and Cas9 were combined and incubated at room temperature for 10 minutes before being combined with the MC38 cells for electroporation. For each electroporation reaction of 200,000 cells with the Neon 10 ul tip, we used 7.5 pmol sgRNA and 7.5 pmol of Cas9. We had previously optimized the electroporation settings to achieve the highest efficiency and cell viability in MC38 cells, which resulted in a voltage of 1200, 30 ms width, and 1 pulse. The MC38 cells were electroporated on the Neon electroporation device (Thermo Fisher). We tested 6 different sgRNAs separately for each allele and filtered out any that had off-target effects on the non-target allele (results shown in Figure S1). We combined the sequences for each allele that did not have an off-target effect to achieve a better knock out. The successful set of sequences that we used for each allele is highlighted in Table S1 in supplemental data.

## Acknowledgements

We thank Paulos Lab members for all the helpful discussions and feedback on this project. This work is supported in part by the Flow Cytometry and Cell Sorting Unit Shared Resource and in part by the Translational Science Translational Science Laboratory at Hollings Cancer Center, Medical University of South Carolina (P30 CA138313). The authors declare no competing financial interests. Finally, we would like to thank Alex Rubinsteyn, Geddes Levenson, Hannah Hendrix, Alper Tunga Ozdemir, Benjamin Vincent for their help in naming MC38 TCRs.

