## Supplemental Data for "Discovering tumor-reactive T-cell receptors through single-cell sequencing of tumor-infiltrating lymphocytes"

| Target | sgRNA sequences (5' to 3') |
|---|---|
| H2D1 1 | ggccccgacucagacccgcg |
| H2D1 2 | cgacgcaagugggagcagag |
| H2D1 3 | uagccgacagagauguaccg |
| H2D1 4 | gugagccugaggaaccugcu |
| H2D1 5 | ugacuucaccuuuagaucug |
| H2D1 6 | agaugucuggcugugacuug |
| H2K1 1 | uagccgacuuccauguaccg |
| H2K1 2 | ggcuccgacucagacccgcg |
| H2K1 3 | uaccagcaguacgccuacga |
| H2K1 4 | caaugagcagaguuuccgag |
| H2K1 5 | ucugcugugaugggucacau |

**Table S1. sgRNA sequences tested to knock out H2-D$^b$ and H2-K$^b$ from the MC38 cell line.** The sgRNA sequences were generated with the online tool by Synthego.

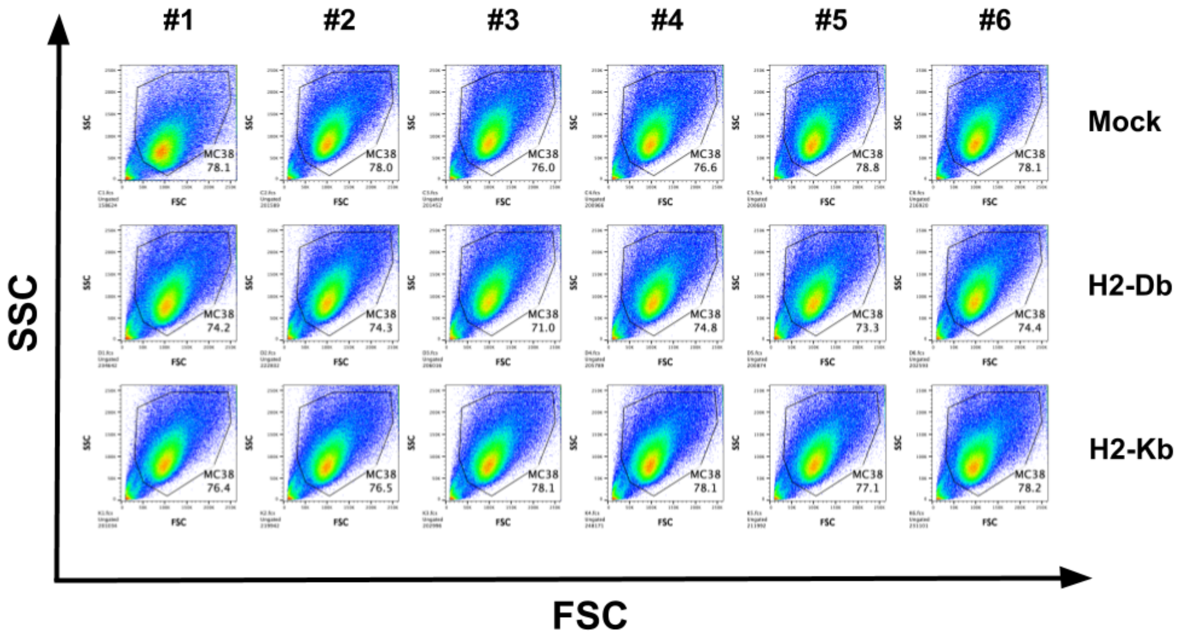

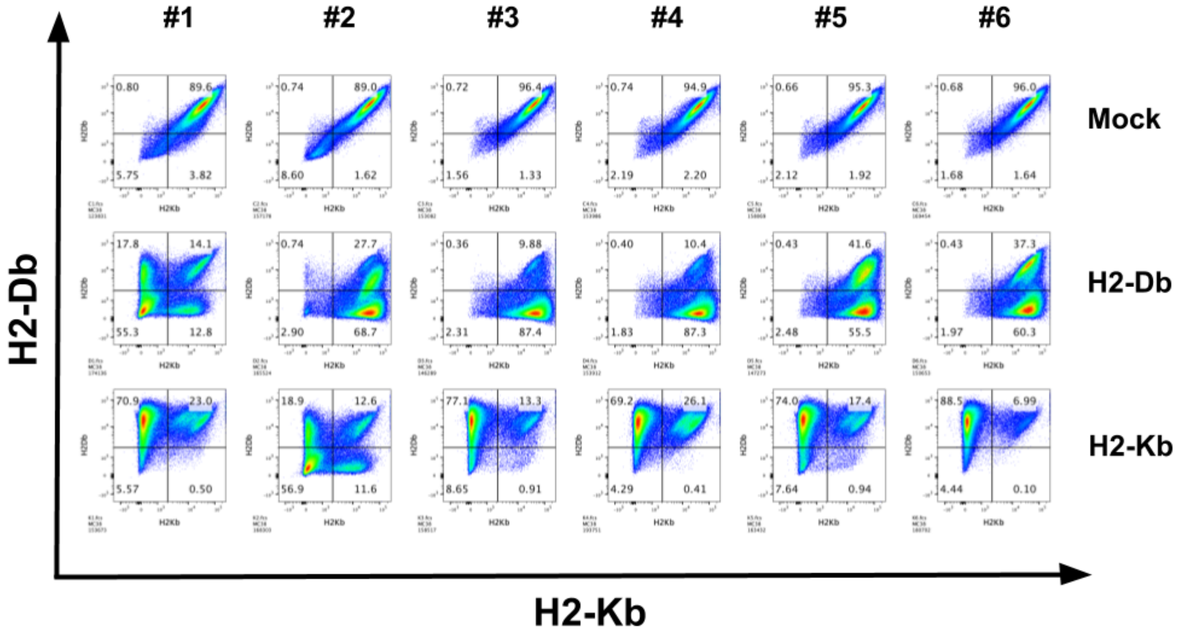


**Figure S1. Six different sgRNAs were tested via electroporation to knock out H2-K$^b$ or H2-D$^b$ in MC38 cells.** Upper panel shows the FSC and SCC for each sample. There was no significant cell death. Lower panel shows H2-K$^b$ expression (x-axis) and H2-D$^b$ expression (y-axis) for each condition.

| Peptide # | Gene | Amino Acid Sequence | Affinity | Peptide length | Solvent |
|---|---|---|---|---|---|
| 1 | Vmn1r88 | YTFVFKPHL | 19.930986 | 9 | Ultrapure water |
| 2 | Hdgfl2 | KGYPHWPAL | 21.3866109 | 9 | Ultrapure water |
| 3 | Zbtb40 | KSFHFYCPL | 22.3497455 | 9 | Ultrapure water |
| 4 | Kmt2b | SNFHFMCAL | 24.5518643 | 9 | Formate acid |
| 5 | Flrt2 | CTFSHLTKL | 26.4282343 | 9 | Ultrapure water |
| 6 | Herc6 | CVYEHTAVL | 26.8884427 | 9 | N-methyl-2-pyrrolidone (Analytical grade) |
| 7 | Copb2 | MSYFLQGTL | 29.5288377 | 9 | N-methyl-2-pyrrolidone (Analytical grade) |
| 8 | Prkd2 | VTYFVGETL | 30.72431 | 9 | DMSO |
| 9 | Spire1 | RSYQYVMKI | 31.9720736 | 9 | Ultrapure water |
| 10 | Ell2 | KAYTKPELL | 32.0075849 | 9 | Ultrapure water |
| 11 | Fn1 | HLYPHVPAL | 32.2305835 | 9 | Ultrapure water |
| 12 | Dvl1 | SSVPVAPQL | 33.7290182 | 9 | Ultrapure water |
| 13 | Syde1 | SAVDFKLHI | 33.7826952 | 9 | Ultrapure water |
| 14 | Elp1 | SSLKSYVQL | 33.862114 | 9 | Ultrapure water |
| 15 | Zc3h7b | VIFTFLCEI | 34.3985341 | 9 | Formate acid |

| | | | | | |
|---|---|---|---|---|---|
| 16 | Huwe1 | ASYNGFLPV | 35.5245222 | 9 | Ultrapure water |
| 17 | Zbtb8os | SAMAMFGYM | 36.7665344 | 9 | Ultrapure water |
| 18 | Man2c1 | SSFLTPEKL | 36.8706879 | 9 | Ultrapure water |
| 19 | Sart3 | SVTVFVNNL | 37.1602419 | 9 | Formate acid |
| 20 | Wdr74 | VSPQHRPVL | 37.5102274 | 9 | Ultrapure water |
| 21 | Ipo11 | SFYELLPEM | 38.3267483 | 9 | N-methyl-2-pyrrolidone (Analytical grade) |
| 22 | Zbtb24 | SLLEHMSLL | 39.0086243 | 9 | Ultrapure water |
| 23 | Dpagt1 | ASIIVFNLL | 39.1469761 | 9 | Formate acid |
| 24 | Chordc1 | KAYQGLQSL | 39.2054976 | 9 | Ultrapure water |
| 25 | Rgl3 | ISPTVRATL | 39.4299159 | 9 | Ultrapure water |
| 26 | Acvr1 | SIVSGLAHL | 44.5017214 | 9 | Ultrapure water |
| 27 | Top2b | VVHQVVSKL | 44.7044938 | 9 | Ultrapure water |
| 28 | Nepro | VLYSTHNRL | 44.7057768 | 9 | Ultrapure water |
| 29 | Wdr12 | ASLDHTIRV | 45.7689268 | 9 | Ultrapure water |
| 30 | Abcb1b | LVYAWQLTL | 45.829361 | 9 | Ultrapure water |
| 31 | Adpgk | ASMTNMELM | 46.6577921 | 9 | 3% ammonia water |
| 32 | Katnb1 | TVLPQIQKL | 46.6767316 | 9 | Ultrapure water |
| 33 | Zfp365 | VAVEIIAVL | 47.2451539 | 9 | 3% ammonia water |
| 34 | Hacd2 | SYIPLFPHL | 47.4071229 | 9 | Ultrapure water |
| 35 | Hace1 | QIYAFLQGF | 48.2310455 | 9 | DMSO |
| 36 | Tbc1d8b | YSFLLGSEL | 48.4042291 | 9 | DMSO |
| 37 | Ttc13 | LTFFHSGLL | 49.3254385 | 9 | Ultrapure water |
| 38 | Cplane1 | QQVNFLSLM | 51.1982012 | 9 | Ultrapure water |
| 39 | Wdr26 | ATPELASSL | 51.5068706 | 9 | Ultrapure water |
| 40 | Ankrd13a | MALEGLPEL | 51.7674559 | 9 | Ultrapure water |
| 41 | Kdm6b | TAPAGTPHL | 51.7877514 | 9 | Ultrapure water |
| 42 | Tor1aip2 | SVAGFNPAL | 51.9957591 | 9 | Ultrapure water |
| 43 | Nphp3 | VVPAPIPPL | 52.0707852 | 9 | Ultrapure water |
| 44 | Pigk | VTVEKFLRV | 52.1143007 | 9 | Ultrapure water |
| 45 | Cpne1 | SSPYSLHYL | 52.3482319 | 9 | Ultrapure water |
| 46 | Gtf3c1 | DVYPFHMIL | 52.5595991 | 9 | Ultrapure water |
| 47 | Ccr10 | AAFLFLACI | 53.8426095 | 9 | Formate acid |
| 48 | Flii | TSLECLSNL | 53.9001703 | 9 | N-methyl-2-pyrrolidone (Analytical grade) |
| 49 | Apeh | VVFDSVQRI | 54.4735558 | 9 | Ultrapure water |

| | | | | | |
|---|---|---|---|---|---|
| 50 | Ulk2 | AAPLLAASL | 55.9631405 | 9 | Ultrapure water |
| 51 | Rev3l | AQHVFQVSL | 56.2936464 | 9 | Ultrapure water |
| 52 | Ltbp3 | VAPGPSARL | 57.2852201 | 9 | Ultrapure water |
| 53 | Grk2 | IVHGYMSKI | 57.749133 | 9 | Ultrapure water |
| 54 | Mfsd2a | CAMGFFLQI | 57.7839656 | 9 | Ultrapure water |
| 55 | Klhl30 | YVYDPGGNL | 58.5472656 | 9 | Ultrapure water |
| 56 | Ampd2 | CVVPFTDLL | 58.6621161 | 9 | Ultrapure water |
| 57 | Lrch3 | AAVPGVLSL | 59.9714954 | 9 | Ultrapure water |
| 58 | Notch2 | VSIPTIAAV | 60.3143185 | 9 | Ultrapure water |
| 59 | Atp13a2 | VVSPFTSCM | 60.5938576 | 9 | Ultrapure water |
| 60 | Tmem198b | SAWVPFGGL | 61.7694108 | 9 | Ultrapure water |
| 61 | Tcf20 | ASYHNMGDL | 61.95914 | 9 | Ultrapure water |
| 62 | Zfp759 | LMLENYNNL | 61.9893876 | 9 | Ultrapure water |
| 63 | Mroh2a | AAVDTLMTL | 62.8208256 | 9 | N-methyl-2-pyrrolidone (Analytical grade) |
| 64 | Psmd2 | VLYGLIAAM | 63.3364415 | 9 | N-methyl-2-pyrrolidone (Analytical grade) |
| 65 | Efnb1 | AAALWLSTL | 63.4392082 | 9 | N-methyl-2-pyrrolidone (Analytical grade) |
| 66 | Irgq | AALLNSAVL | 63.8404242 | 9 | Ultrapure water |
| 67 | Xab2 | QLYELALKL | 64.1369137 | 9 | Ultrapure water |
| 68 | Med20 | AVYGPSDTM | 65.2400578 | 9 | Ultrapure water |
| 69 | Apex2 | TSVVELPTL | 65.3124145 | 9 | Ultrapure water |
| 70 | Sesn2 | RAPEKLLKL | 65.8819993 | 9 | Ultrapure water |
| 71 | Yipf4 | VAYGQVLAV | 66.2200647 | 9 | Ultrapure water |
| 72 | Vcam1 | AVFGASTVL | 67.2668472 | 9 | DMSO |
| 73 | Dyrk1b | KARKYFELL | 68.1329769 | 9 | Ultrapure water |
| 74 | Gm49336 | VIFQDVENL | 68.7738695 | 9 | DMSO |
| 75 | Prpsap1 | AGLTHIITI | 68.9834763 | 9 | Ultrapure water |
| 76 | Rhd | VNLVQLTLM | 69.6456672 | 9 | Ultrapure water |
| 77 | Aatf | MAPIDHTTM | 71.1486327 | 9 | Ultrapure water |
| 78 | N4bp2l2 | VATINFRRL | 71.7568319 | 9 | Ultrapure water |
| 79 | Zfp775 | KSFRQKPNL | 71.9107416 | 9 | Ultrapure water |
| 80 | Dpagt1 | IIVFNLLEL | 73.3660407 | 9 | 3% ammonia water |
| 81 | Ksr1 | LAPEIVLEM | 73.9692439 | 9 | Ultrapure water |
| 82 | Med1 | FSLSFQHPV | 74.8922638 | 9 | Ultrapure water |

| | | | | | |
|---|---|---|---|---|---|
| 83 | Zfyve26 | LSYQRVSGV | 75.0085607 | 9 | Ultrapure water |
| 84 | Aoc2 | IVGHFYGGL | 75.4376487 | 9 | Ultrapure water |
| 85 | Trmt1l | VSMPGKTAI | 75.4461776 | 9 | Ultrapure water |
| 86 | Gm6502 | MNVHTPPKL | 75.4829802 | 9 | Ultrapure water |
| 87 | Bbs10 | SGVEVFEFL | 76.2545262 | 9 | Ultrapure water |
| 88 | Ptprs | TAVVRQSTL | 76.50261 | 9 | Ultrapure water |
| 89 | Abcb1b | LSLVYAWQL | 76.9261987 | 9 | N-methyl-2-pyrrolidone (Analytical grade) |
| 90 | Cd81 | AIQEFQCLL | 77.4249794 | 9 | N-methyl-2-pyrrolidone (Analytical grade) |
| 91 | Nbeal2 | VLIQWLPAL | 78.3292866 | 9 | N-methyl-2-pyrrolidone (Analytical grade) |
| 92 | Mfsd2a | MGFFLQIYL | 78.9389669 | 9 | Formate acid |
| 93 | Notch2 | VTFQDAANM | 79.2454181 | 9 | 3% ammonia water |
| 94 | Slc23a2 | ITVVGLSNL | 79.7459564 | 9 | Ultrapure water |
| 95 | Eipr1 | ATLEGRGHL | 80.8010109 | 9 | Ultrapure water |
| 96 | Ccr10 | YSASFHAAF | 81.3284452 | 9 | Ultrapure water |

**Table S2. Predicted MC38 Neoantigen peptides used in the peptide screen.** The gene from which the peptide was derived, the amino acid sequence, predicted affinity, length of peptide, and solvent used for reconstitution are listed.
